# Predicting cell stress and strain during extrusion bioprinting

**DOI:** 10.1101/2022.09.28.509836

**Authors:** Sebastian J. Müller, Ben Fabry, Stephan Gekle

## Abstract

Bioprinting of living cells can cause major shape deformations, which may severely affect cell survival and functionality. While the shear stresses occurring during cell flow through the printer nozzle have been quantified to some extent, the extensional stresses occurring as cells leave the nozzle into the free printing strand have been mostly ignored. Here we use Lattice-Boltzmann simulations together with a finite-element based cell model to study cell deformation at the nozzle exit. Our simulation results are in good qualitative agreement with experimental microscopy images. We show that for cells flowing in the center of the nozzle extensional stresses can be significant, while for cells flowing off-center their deformation is dominated by the shear flow inside the nozzle. From the results of these simulations, we develop two simple methods that only require the printing parameters (nozzle diameter, flow rate, bioink rheology) to (i) accurately predict the maximum cell stress occurring during the 3D bioprinting process and (ii) approximately predict the cell strains caused by the elongational flow at the nozzle exit.

## I. INTRODUCTION

The aim of 3D bioprinting is to transfer the well-established techniques of conventional 3D printing to the fabrication of functional, living tissues. The material to be printed typically consists of a chemically complex hydrogel, termed the bioink, in which living cells are suspended. This technology promises to become a major breakthrough, e.g. for cancer studies or – in the long run – organ replacements [1–5]. A key obstacle, however, remains to ensure the survival and functionality of cells during and after the fabrication process. Possible causes for cell damage are numerous, but can be broadly classified into insufficient bio-compatibility and mechanical damage. The former arises from direct interaction between the cell and the surrounding bioink and has been intensively studied [6–13].

Mechanical damage, by contrast, arises from hydrodynamic stresses as the cell passes from the reservoir through the printing nozzle, transitions into the printing strand, and finally comes to rest in the printed construct. It has been shown that even after optimizing biological and chemical conditions [14], such hydrodynamic stresses remain a crucial source of cell damage and death [15–25]. Understanding these mechanical stress response processes is notoriously difficult as they result from an interplay between the complex rheology of the bioink, which is typically shear thinning [26–29], and the even more complex viscoelastic response of the cell itself [30–39]. Despite these difficulties, certain progress towards reliable theoretical estimates of the cell stress inside printing needles has been achieved [15, 18, 40]. As a starting point, the fluid shear stress profiles within printing nozzles have been computed theoretically [17, 21, 41]. Some experimental studies correlated such stress calculations with cell viability or functionality [15, 22, 40, 42–47]. These studies, however, investigated hydrodynamic stresses only up to the end of the printing nozzle. At the transition from the nozzle exit into the free strand, the flow profile transitions rapidly from a Poiseuille-like profile in the nozzle to a plug flow profile inside the free bioink strand. This transition is accompanied by sizable radial flows whose effect on cell deformation and therefore cell damage has so far not been experimentally or theoretically quantified.

In this work, we start with fully three-dimensional Lattice Boltzmann calculations for the flow profile of shear thinning fluids [41] at the exit of a printing nozzle. To investigate cell stress and strains inside and during exit from the printing nozzle, we then employ our recently developed hyperelastic cell model [30] which includes explicit two-way coupling between bioink and cellular mechanics, and show its qualitative match with experimental micrographs taken during the printing process. From these investigations, we finally develop simple methods to predict the cell stress and strains occurring during bio-printing processes, and specifically during nozzle exit, by only using the printing parameters, i. e., the nozzle diameter, the bioink rheology, and the volumetric flow rate, as input quantities. For this, we combine the classical theories of Jeffery [48] and Roscoe [49] with our semi-analytical flow computations of shear thinning bioinks in capillaries [41].

## II. METHODS AND SETUP

### A. Flow dynamics: Lattice-Boltzmann simulations

In our simulations, we employ a fully three-dimensional fluid dynamics solver. We use the implementation of the Lattice Boltzmann method [50] provided by the software package ESPResSo [51, 52], which we extended with algorithms to allow for the simulation of free-slip surfaces [53] and shear thinning fluids [41]. Using an immersed-boundary algorithm [32, 54–56], we couple our cell model (section II C) to the flow.

### B. Bioink rheology

The shear thinning rheology is considered an essential material property for bioinks, as it serves two purposes: first, the large viscosity of the material at rest provides the necessary mechanical stability of the printed construct itself. Second, the shear thinning properties allow the material to be printed at significantly lower pressure — considering the same printing speed —, thus reducing the overall hydrodynamic stresses acting on cells inside the nozzle.

We describe the viscosity as function of the rate of strain 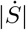 according to a three-parameter simplified Carreau-Yasuda model, also known as Cross model [41, 57]:

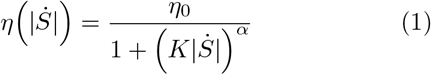

Here, *η*_0_ is the zero-shear viscosity and the exponent *α* characterizes the shear thinning strength of the bioink, with *α* = 0 for a Newtonian fluid, and *α >* 0 for a shear thinning fluid. The inverse of the time constant, *K*^−1^, defines the rate of strain at which the viscosity is equal to *η*_0_*/*2. 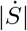 is calculated as the contraction of the rate of strain tensor 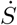 via

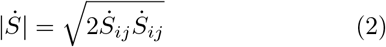

with the tensor elements

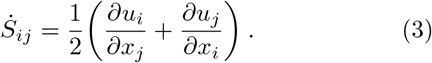

The diagonal elements of 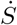 are the rates of elongation of the fluid along the coordinate axes, and the off-diagonal elements are the respective shear rates. We choose *η*_0_ = 10 Pa s and *K* = 0.05 s for the zero-shear viscosity and the time constant, respectively. This parameter choice roughly resembles the values obtained for 2.5% alginate hydrogels [26, 41, 58] which is a widely used bioink material. In order to investigate the influence of the shear thinning strength in our calculations, we pick six different values for *α* between 0 and 1 with *α* = 0.75 [41] corresponding to the said alginate solution. The viscosity as function of the shear rate is depicted in figure 1(a). For an idealized, i.e. infinitely long, cylindrical nozzle, the velocity profile and the fluid stress *σ*_f_ can be computed according to [41][59] as shown in figure 1(b and c) with the pressure adjusted so as to ensure the same flow rate for each *α*. In our previous study [41], we introduced this method to calculate the velocity, shear rate, viscosity, and shear stress, profiles for an inelastic shear thinning fluid in a cylindrical nozzle. The central assumptions — a laminar, uni-axial, pressure driven, flow — are usually applicable for the description of bioink extrusion. In the following, we define the fluid stress as:

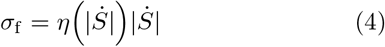

**FIG. 1.**
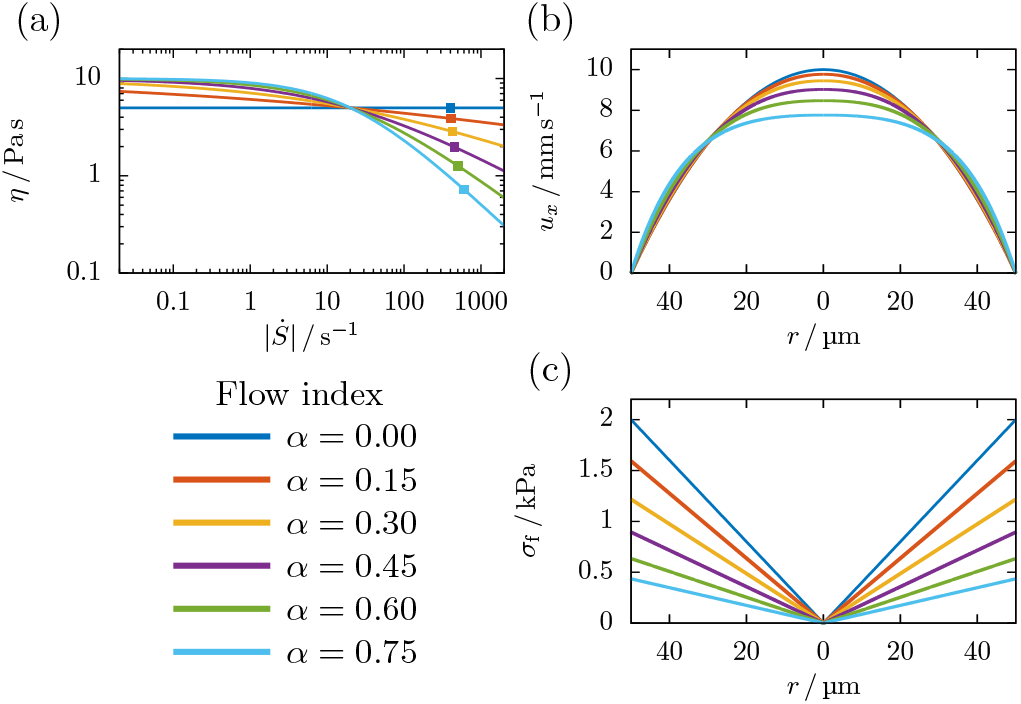
(a) Viscosity as function of the shear rate for the six different degrees of shear thinning (*α* = 0, 0.15, 0.3, 0.45, 0.6,and 0.75). Squares indicate the maximum shear rate in the nozzle channel under the flow conditions used in our simulations (cf. section II D). (b) Corresponding velocity profiles for an average velocity of 5 mm s^−1^ inside the cylindrical printing nozzle. With increasing shear thinning strength, the velocity profile flattens at the center. (c) Corresponding fluid stress profiles. Stresses are linear with the radial position, and the maximum fluid stress decreases significantly with increasing *α* at constant flow rate.

We note that, if a constant extrusion pressure was used for calculation, the fluid stress profile in figure 1(c) would be the same regardless of *α* [41].

### C. Cell elasticity

#### 1. Hyperelastic cell model

Our cell is modeled as hyperelastic continuum, with a sphere as equilibrium configuration. We provide extensive validation of the model in a previous publication [30]. This includes AFM and FluidFM® measurements on biological cells and hydrogel particles as well as comparison to analytical theories [38, 49] and previous numerical simulations in shear flow [32, 60].

As a hyperelastic model, we employ the neo-Hookean material model. This model is strain-hardening for compressive strain, e.g. in AFM experiments, but also for shear strains as occurring mainly in microfluidic experiments. Its strain energy density is computed via [61, p. 100]

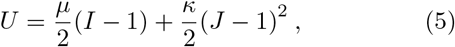

where *J* = det(*F*) is the determinant of the deformation gradient tensor 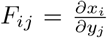 [61, p. 14, 18], with the undeformed and deformed vertex coordinates *x*_*i*_ and *y*_*i*_, respectively. *I* = *J* ^−2*/*3^ tr(*F* ^**T**^*F*) denotes the second invariant of *F*. As elastic parameters we choose a shear modulus of *µ* = 1000 Pa and a Poisson ratio of *ν* = 0.48. A simulation series with *µ* = 500 Pa is included in section S-9 of the Supplementary Material. The Poisson ratio near 0.5 provides sufficient incompressibility of the cell, while the shear modulus lies in the range typically found for mammalian stem cells [62]. In consistency with linear elasticity for small deformations, the shear and bulk modulus relate to the Young’s modulus and Poisson ratio via

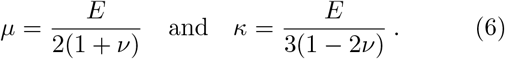

The cell radius is chosen as *R*_c_ = 8 µm (6 in simulation units), and the mesh consists of 9376 tetrahedra.

In our numerical method, the interior of the cell is filled with the same fluid as the outside fluid. Together with the Neo-Hookean elasticity, this leads to an effectively viscoelastic cell response [30].

#### 2. Force calculation and flow coupling

For numerical simulations, the spherical volume is uniformly tetrahedralized using the meshing software Gmsh [63]. The elastic forces acting on each vertex of one tetrahedron are obtained via differentiation of the strain energy density (5) with respect to the relative vertex displacement,

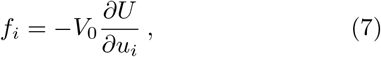

where *V*_0_ denotes the reference volume of the tetrahedron and *u*_*i*_ = *y*_*i*_ − *x*_*i*_. This approach is explained in detail in section 3.1 in [30].

The coupling between the cell model and the bioink is realized using an immersed-boundary algorithm [64, 65]. After computation of the cell vertex forces, they are transmitted into the fluid via extrapolation from the moving Lagrangian cell mesh onto the static Eulerian Lattice Boltzmann grid. The two-way coupling is completed through advecting the cell vertices with the local interpolated fluid velocity.

#### 3. Cell stress calculations

In addition to the elastic forces, we are able to obtain the internal stress distribution inside our cell model. We compute the Cauchy stress tensor in each tetrahedron from the strain energy density and the deformation gradient tensor according to Bower [61, p. 97] as:

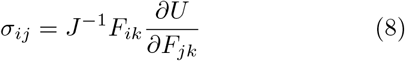

For the neo-Hookean model in (5), this becomes

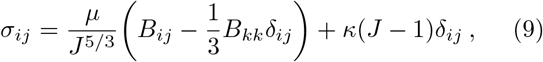

where *B* = *FF* ^**T**^ denotes the left Cauchy-Green deformation tensor.

In order to obtain a simple scalar observable to quantify the cell stress, we start from the local von Mises stress in each tetrahedron. The von Mises stress is an invariant of the Cauchy stress tensor and commonly used in plasticity theory to predict yielding of materials under multiaxial loading conditions through construction of a fictitious uniaxial loading. Using the principal stresses, i. e., the eigenvalues *σ*_1_, *σ*_2_, and *σ*_3_, of the Cauchy stress tensor (9), we calculate [61, p. 48]

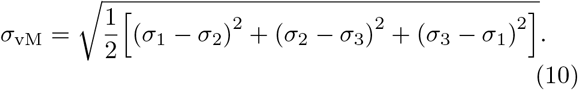

An alternative equivalent formulation to (10) is the contraction of the deviator of the Cauchy stress tensor 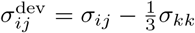. It reads:

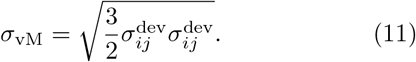

The total cell stress *σ*_vM_ is then computed by averaging the local von Mises stress over all tetrahedra in the cell model weighted by the undeformed volume of each tetrahedron.

#### 4. Validation of the cell stress calculation

We validate our cell stress calculations using a linear shear flow setup: the simulation box with dimensions 10 × 15 × 5 (*x* × *y* × *z* in units of *R*_c_) is bounded by two planes at *y* = 0 and *y* = 15*R*_c_, moving with a tangential velocity in ±*x*-direction. This creates a linearly increasing velocity across the gap and thus a constant shear rate 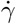 in the box. The shear rate is varied to achieve a range of fluid stresses up to 1.5 kPa, while the fluid viscosity (*α* = 0) and the cell’s shear modulus remain constant. In non-dimensional terms, this range corresponds to capillary numbers 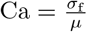 between 0 and 1.5.

During the simulation, the initially spherical cell traverses through a series of ellipsoidal deformations before reaching a stationary state, at which the whole cell volume performs a tank-treading motion, i. e., the cell vertices rotate around the fixed ellipsoidal cell shape. In figure 2, we compare the elastic cell stress in the stationary state calculated by (10) to the analytical calculations of Roscoe [49] (detailed in section S-4) and find excellent agreement for a realistic range of fluid stresses.

**FIG. 2.**
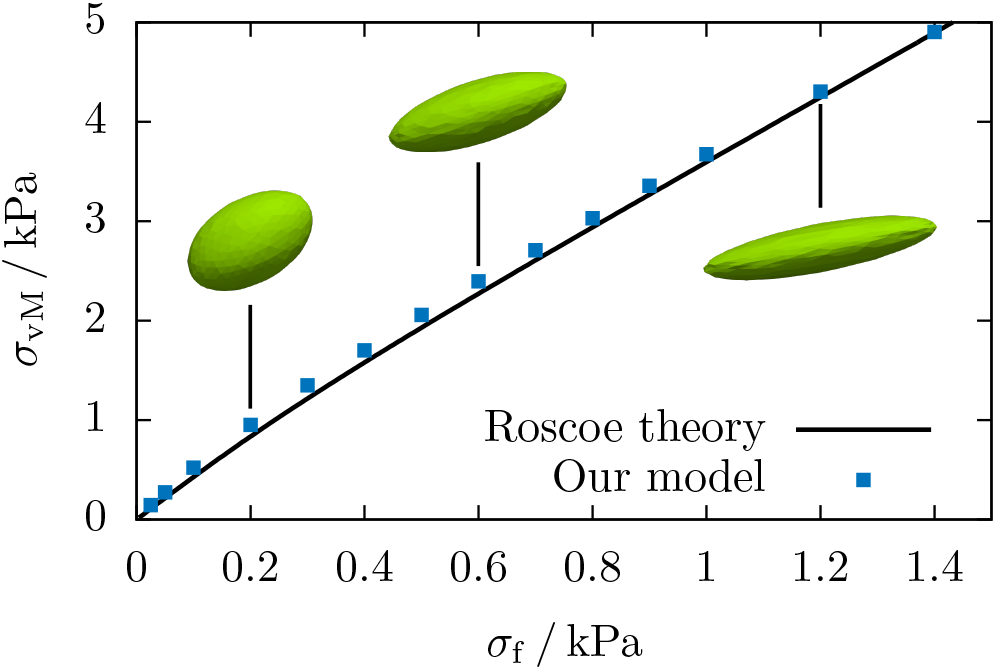
Comparison of the cell stress predicted by Roscoe [49] and the average cell stress of our model in shear flow. Insets show the stationary, tank-treading shape of the simulated cell at fluid stresses corresponding to Ca = 0.2, 0.6, and 1.2.

In addition to the elastic stress, we compute the viscous contribution resulting from the fluid motion enclosed by the cell volume. This quantity is extracted from the Lattice-Boltzmann strain rate tensor field [41, 66] inside the cell using our method from [67] and averaging over the cell volume. In figure S-2 we show that the agreement of the numerically obtained viscous cell stress with Roscoe theory is equally good as for the elastic component.

We note that cell and fluid stress in figure 2 are time-independent and stationary. We further dissect their relation in detail in section III A.

### D. Bioprinting simulations

The two essential parts of the bioprinting process are (i) the flow inside the nozzle channel and (ii) the flow transition at the nozzle exit. Both situations will be studied separately in this work.

#### 1. Nozzle channel

We model the nozzle channel using a periodic cylindrical no-slip channel with a radius of *R* = 50 µm and length of 133 µm (37.5 and 100 in simulation units), as depicted in the left dashed box in figure 3. The shear thinning fluid dynamics are solved by the Lattice-Boltzmann method as described in section II A. No-slip boundary conditions are imposed at the channel wall. The flow is driven by a pressure gradient *G* along the nozzle axis. To compare the different bioinks detailed in section II B, we consider a fixed average velocity of 5 mm s^−1^ (volumetric flow rate Ω = 3.93 × 10^−11^ m^3^ s^−1^ ≈141 µl h^−1^). The corresponding pressure gradient is different for each *α* and is obtained according to [41]. Our input parameters as well as averaged and maximum quantities of the nozzle channel flow are listed in table I. We note that compared to common flow cytometry setups [31, 35], the channel radius in typical bioprinting applications is significantly larger, thus allowing cells to flow off-centered. To account for this, a single spherical cell is inserted at different radial starting positions of 0, 1.5 *R*_c_, 3 *R*_c_, and 4.5 *R*_c_, as shown in figure 3.

**FIG. 3.**
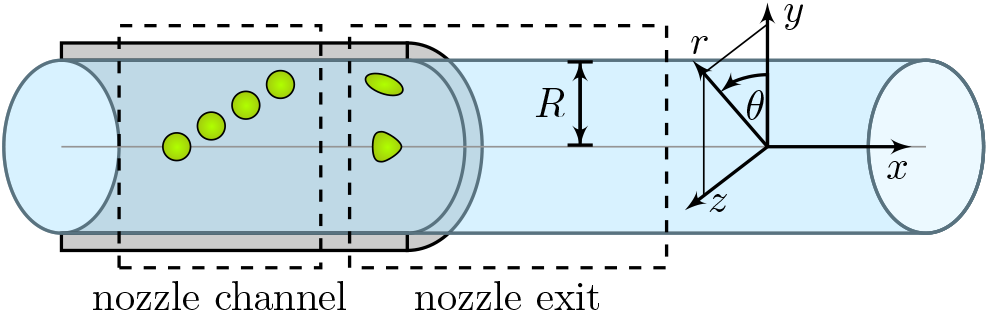
Schematic of the bioprinter setup: the nozzle channel is bounded by a cylindrical wall and periodic in *x*-direction. Single cells are inserted at different radial offsets. The nozzle exit consists of the transition from the no-slip nozzle channel to the free bioink strand. Cells are initialized in a deformed state close to the transition.

**TABLE I.** Flow parameters of our bioprinter setup with a nozzle radius of 50 µm and an average velocity of 5 mm s^−1^.

| $\alpha$ | $G$<br>$\text{Pa m}^{-1}$ | $\sigma_f^{\text{max}}$<br>kPa | $\sigma_f^{\text{avg}}$<br>kPa | $u_x^{\text{max}}$<br>$\text{mm s}^{-1}$ | $\vert \dot{S}\vert ^{\text{max}}$<br>$\text{s}^{-1}$ | $\vert \dot{S}\vert ^{\text{avg}}$<br>$\text{s}^{-1}$ |
| --- | --- | --- | --- | --- | --- | --- |
| 0.00 | $8.00 \times 10^7$ | 2.00 | 1.33 | 10.0 | 400 | 267 |
| 0.15 | $6.37 \times 10^7$ | 1.59 | 1.06 | 9.77 | 410 | 265 |
| 0.30 | $4.87 \times 10^7$ | 1.22 | 0.81 | 9.45 | 426 | 262 |
| 0.45 | $3.58 \times 10^7$ | 0.89 | 0.60 | 9.03 | 454 | 257 |
| 0.60 | $2.54 \times 10^7$ | 0.63 | 0.42 | 8.47 | 502 | 251 |
| 0.75 | $1.74 \times 10^7$ | 0.44 | 0.29 | 7.77 | 604 | 242 |

#### 2. Nozzle exit

The geometry of our simulations at the nozzle exit is depicted by the right dashed box in figure 3, the flow dynamics are again solved by the Lattice-Boltzmann method. The free bioink strand of length 933 µm (700 in simulation units) behind the nozzle exit is assumed to have the same radius as the inner radius of the nozzle channel, with free-slip boundary conditions applied at the fluid surface, which result in a plug motion of the bioink. This way we neglect the small extension of the bioink strand at the nozzle exit known as Barus effect or die swell [68]. Equal flow conditions as inside the nozzle channel are achieved by applying the average velocity of 5 mm s^−1^ as normal velocity at the circular inflow and outflow planes, instead of a constant pressure gradient as used in the nozzle channel setup. We insert a single cell at different radial positions as explained above. The starting configuration of the cell is taken from the corresponding simulation of the nozzle channel setup, i. e., the cell is inserted in a deformed state close to the nozzle exit, as shown in the first frames in figure 9.

## III. RESULTS

### A. Dissecting the notion of “cell stress”

In many situations, it has become a common approach to invoke the term “cell stress” and to equate it directly to the fluid stress, i. e., the viscosity multiplied by the local shear rate at the cell position. Here, we show that this simple approach, while being correct in its order of magnitude, hides a good amount of the more complex features of intracellular stress. To illustrate this, we apply the theory of Roscoe [49] for a cell in linear shear flow, which accurately describes cell behavior in our numerical simulations (see section II C 4) and in microchannel experiments [69], provided that the cell does not flow in the channel center where the shear rate approaches zero.

Inside a flowing cell, two qualitatively different kinds of stress arise. The first kind are viscous stresses that are caused by frictional motion (i. e. tank-treading) of the cell interior. The second kind are elastic stresses that are caused by the deformation (e. g. shearing and stretching) of the cell. The magnitude of the former are governed by the cell’s internal viscosity, while the latter are set by its elastic moduli. We note that, in principle, both a cell’s viscosity and its elasticity can be non-homogeneous, i. e., they vary spatially throughout the cell, and anisotropic, i. e., they depend on direction, e. g., due to alignment of certain cytoskeletal elements. Here and in most other situations, these more complicated effects are neglected, and the cell is considered a homogeneous, isotropic viscoelastic medium. Furthermore, as shown in [49] for a cell in pure shear flow, stability requires that viscous and elastic cellular stresses do not vary between different locations inside the cell. Their value can be calculated from Roscoe theory as detailed in the Supporting Information (eqs. (S-43) and (S-48)).

We start with the limiting case of low shear rates corresponding to small capillary numbers 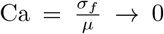. In this limit, fluid stresses are not sufficient to cause significant cell deformation, and the cell essentially remains spherical. Indeed, the classical calculation for a rigid sphere in shear flow detailed in section S-6 of the Supporting Information yields a surprisingly accurate description of this limit. The cell rotates as a rigid body, which implies the absence of internal frictional motions and thus leads to a vanishing viscous cell stress as shown by the purple curve in figure 4. Similarly, elastic stresses in the vorticity direction vanish as shown by the *σ*_33_ component in figure 4. A positive stress appears in a direction inclined by 45° with respect to the flow direction (*σ*_11_), with a corresponding negative stress in the perpendicular direction. Their magnitude is precisely 5*/*2 times the undisturbed fluid stress *σ*_f_, which exactly corresponds to the situation of the rigid sphere as shown in section S-6.

**FIG. 4.**
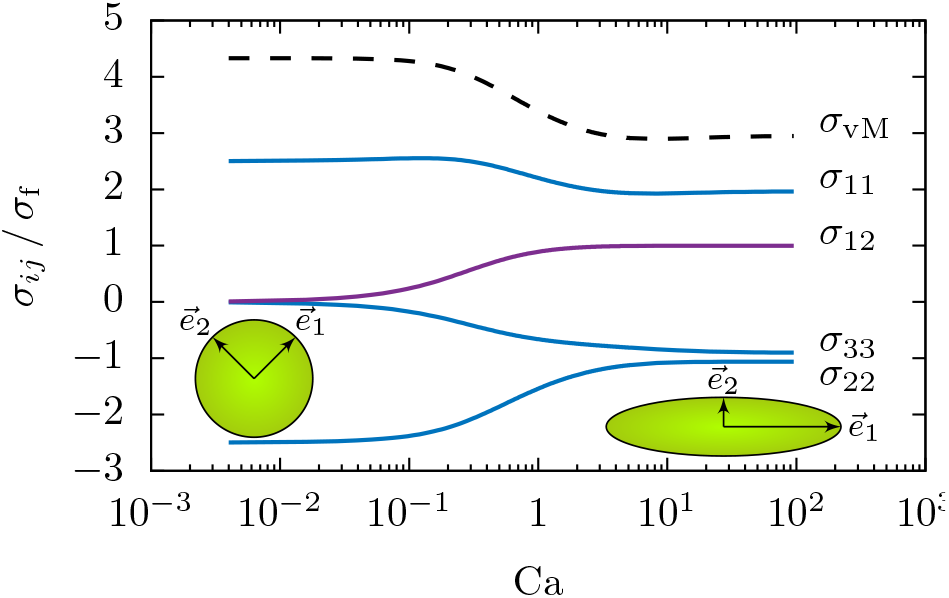
Components of the cell stress tensor *σ*_*ij*_ normalized by the undisturbed fluid stress *σ*_f_ across multiple orders of magnitude of the Capillary number computed using the Roscoe theory. Elastic stresses are shown in blue, viscous stresses are shown in purple. Components not appearing in the figure remain zero throughout. The directions 1, 2, 3 refer to a body-fixed coordinate system as indicated by the insets.

In the opposite limit of high shear rates (Ca ⟶ ∞), the situation becomes more involved. In agreement with our numerical simulations shown in figure 2, the cell becomes strongly elongated and aligned in flow direction. Due to the persisting tank-treading motion, internal viscous stresses do not disappear. Instead, the flatness of the cell shape minimizes the cell’s disturbing influence on the surrounding fluid flow, and indeed the cell’s internal viscous stress now becomes equal to the undisturbed fluid stress, as can be seen by the purple curve in figure 4. Maintaining the flattened cell shape, however, in addition requires elastic stresses. As shown by the blue curve in figure 4, all three elastic stress components arise with their ratios being *σ*_11_ : *σ*_22_ : *σ*_33_ = 2 : −1 : −1. The positive stress in flow direction, *σ*_11_ is balanced by negative stresses in the two other directions. These ratios can easily be understood by the analogy with a uniaxially stretched beam as detailed in section S-7 of the Supporting Information.

Despite this complexity, it may be helpful in many situations to have at hand a single measure to quantify “cell stress”. Such a measure can be provided by the elastic von Mises stress *σ*_vM_ given in (11) which we include as the black dashed line in figure 4. The ratio 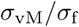 transitions from 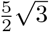 at low to 3 at high Ca. In the intermediate range, the proportionality factor is situated between these two limits. As can also be deduced from figure 2, the relation between *σ*_vM_ and *σ*_f_ changes the most in the range of 0.1 *<* Ca *<* 1, while otherwise an approximately linear dependency emerges.

From the results of this subsection, we conclude that the common approach of equating (undisturbed) fluid stress to “cell stress” can be a reasonable approximation for low and high Capillary numbers.

### B. Cell flowing inside the nozzle channel

Using our setup described in section II D 1, we investigate the cell behavior and the cell’s internal stress distribution during the flow inside the nozzle. Depending on the initial radial position, we observe two modes of deformation of the cell:

i. A cell flowing along the axis of the nozzle channel assumes an axisymmetric bullet-like shape, as can be seen in figure 5(a) and (b) for a Newtonian and a highly shear thinning bioink, respectively. In both cases, the radial dependency of the internal cell stress is highly non-homogeneous and resembles the linearly increasing fluid stress of the undisturbed liquid (cf. figure 1(c)), since the cell’s surface has to balance higher fluid shear stresses for increasing *r* in the stationary state. However, the magnitude of the stress and likewise the cell deformation decrease significantly when the shear thinning index *α* is increased at the same volumetric flow rate. This finding may explain earlier experimental observations in which more shear thinning bioinks were found to increase cell survival in bioprinting [22, 44] when the pressure was reduced to ensure equal flow rates for all conditions.
ii. A cell flowing off-center deforms into an approximately ellipsoidal shape exhibiting tank-treading motion. Due to the curvature of the flow, the cell migrates towards the channel center (sometimes referred to as margination), where it eventually assumes the bullet-like shape discussed in the previous paragraph. A sequence of simulation snapshots for a cell flowing in the Newtonian bioink is shown in figure 5(c), where the internal stress distribution of the off-centered cells can be observed. Figure 6(a) shows the corresponding development of the radial position over time starting from an offset of 36 µm. With increasing shear thinning strength, i. e., decreasing pressure gradient, the cell takes longer to migrate towards the channel center.

In figure 6(b) the same situation is studied for a constant pressure gradient. We find that here the migration speed of the cell becomes independent of the shear thinning properties of the bioink and thus conclude that cell migration is determined predominantly by the applied pressure gradient and not the flow speed. This finding can be understood since the stress, and thus the forces, that the cell feels are directly determined by the local fluid stress. Therefore, when printing bioinks with different rheology at the same printing pressure, the radial cell distribution will not change. When printing bioinks with increasing shear thinning strength at the same flow rate, by contrast, fewer cells will migrate to the center of the nozzle.

**FIG. 5.**
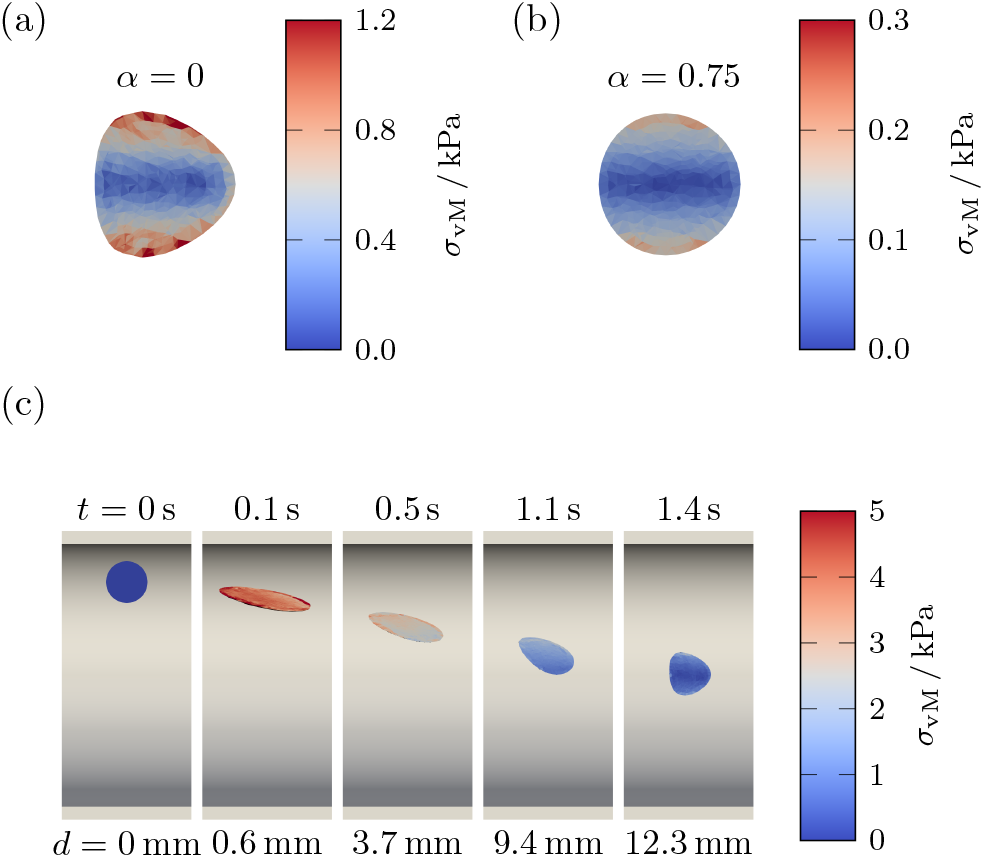
Internal stress distribution for a cell flowing at the center of the nozzle channel in (a) a Newtonian fluid and (b) a strongly shear thinning fluid. (c) Internal stress distribution and radial migration of an off-centered cell towards the axis of the nozzle channel in a Newtonian bioink. The bottom labels give the axial distance traveled during the time given by the top labels.

**FIG. 6.**
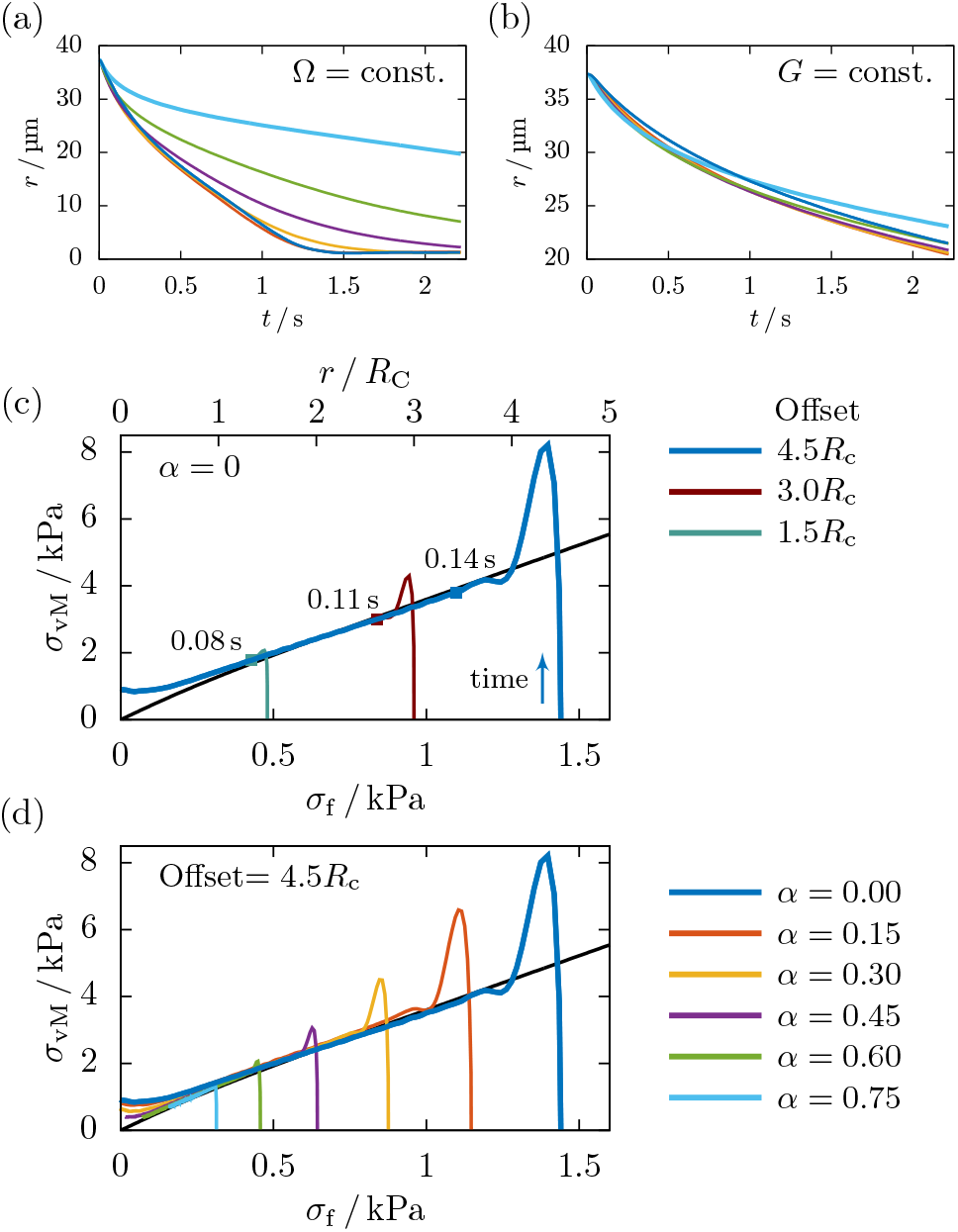
Radial migration *r* (center-of-mass) of a cell starting near the nozzle wall for different shear thinning strengths (see (d) for color labels) with (a) constant flow rate and (b) constant pressure gradient (*G* = 1.14 ×10^7^ Pa m^−1^). For constant *G*, the migration speed is almost independent of *α*. (c,d) Cell stress as function of the local fluid stress compared to the Roscoe theory (black line). Due to the radial migration of the cell, the cell experiences a continuous change of the local fluid stress over time. (c) The cell starting at different offsets (from right to left: 4.5*R*_c_, 3*R*_c_, and 1.5*R*_c_) in the Newtonian fluid. The duration of the deformation from the spherical reference shape to the approximately elliptical shape is given by the points. (d) Cells starting at offset 4.5*R*_c_ for bioinks with increasing shear thinning strength in comparison with the Roscoe theory. For details, see text.

The ellipsoidal cell shape during the migration allows us to compare the cell stress to the prediction of the Roscoe theory [49] detailed in section II C 4. In figure 6(c) we plot the development of the cell stress in a Newtonian bioink when cells are initially placed at different offsets from the nozzle center. Due to the migration of the cells towards the channel center, the local fluid stress experienced by a given cell decreases monotonically over time. In order to directly compare with the prediction of Roscoe theory, which assumes a constant fluid shear stress, we choose this local fluid stress as abscissa. Cells start at offsets of 12 µm, 24 µm and 36 µm corresponding to initial fluid stresses of *σ*_f_ ≈0.5 kPa, 1.0 kPa, and 1.5 kPa, respectively. The initial shape is undeformed and thus *σ*_vM_ = 0 for *t* = 0. The cell first experiences a transient of large stresses and quickly relaxes towards the cell stress predicted by Roscoe where the curved flow is locally approximated as a pure shear flow, as indicated by the square symbols. Due to the migration towards the channel center, the cell stress decreases with time and radial position. The curves of all initial radial offsets perfectly agree with the prediction of the Roscoe theory, as long as the cell’s radial position is larger than *R*_c_. When the cell is close to the channel center, the local shear flow approximation becomes invalid, thus causing deviations from the theoretical prediction.

A similar plot is provided for shear thinning bioinks in figure 6(d) where the stress of cells starting at offset 4.5*R*_c_ for different *α* is compared with Roscoe theory. We again find excellent agreement with the Roscoe theory independent of the shear thinning strength. This finding may seem surprising at first, as the theory of Roscoe is designed for purely Newtonian fluids surrounding the cell, but stays valid for shear thinning bioinks as well. This demonstrates that the key property determining cell motion is indeed not the shear rate, but rather the shear stress. The plots for the remaining cell offsets and bioinks are included in the SI (cf. figure S-3).

### C. Analysis of the flow field at the nozzle exit

In this section, we investigate the influence of the shear thinning rheology of the bioinks introduced in section II B on the undisturbed (cell free) flow field at the nozzle exit, where the transition from nozzle channel to the free bioink strand causes additional radial flows. We use the second setup described in section II D, without a cell, and run the calculations until the flow becomes stationary. In figure 7(a) and (b), we show *x*-*y*-slices of the velocity profiles for the axial and radial velocity, respectively, at different values of the shear thinning parameter *α*. From top to bottom, the shear thinning strength of the fluid increases, while the flow rate is kept constant. The axial velocity component in figure 7(a) shows the same trend for increasing *α* as seen in figure 1(b): the flow develops a central plateau inside the nozzle channel which at the nozzle exit transitions into the plug flow inside the bioink strand. Indeed, as shown in figure 7(c), the ratio 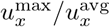 between the maximum velocity inside the nozzle channel and the average velocity assumes the Poiseuille value of 2 at *α* = 0 and decreases towards the plug-flow value of 1 for increasing shear thinning strength.

**FIG. 7.**
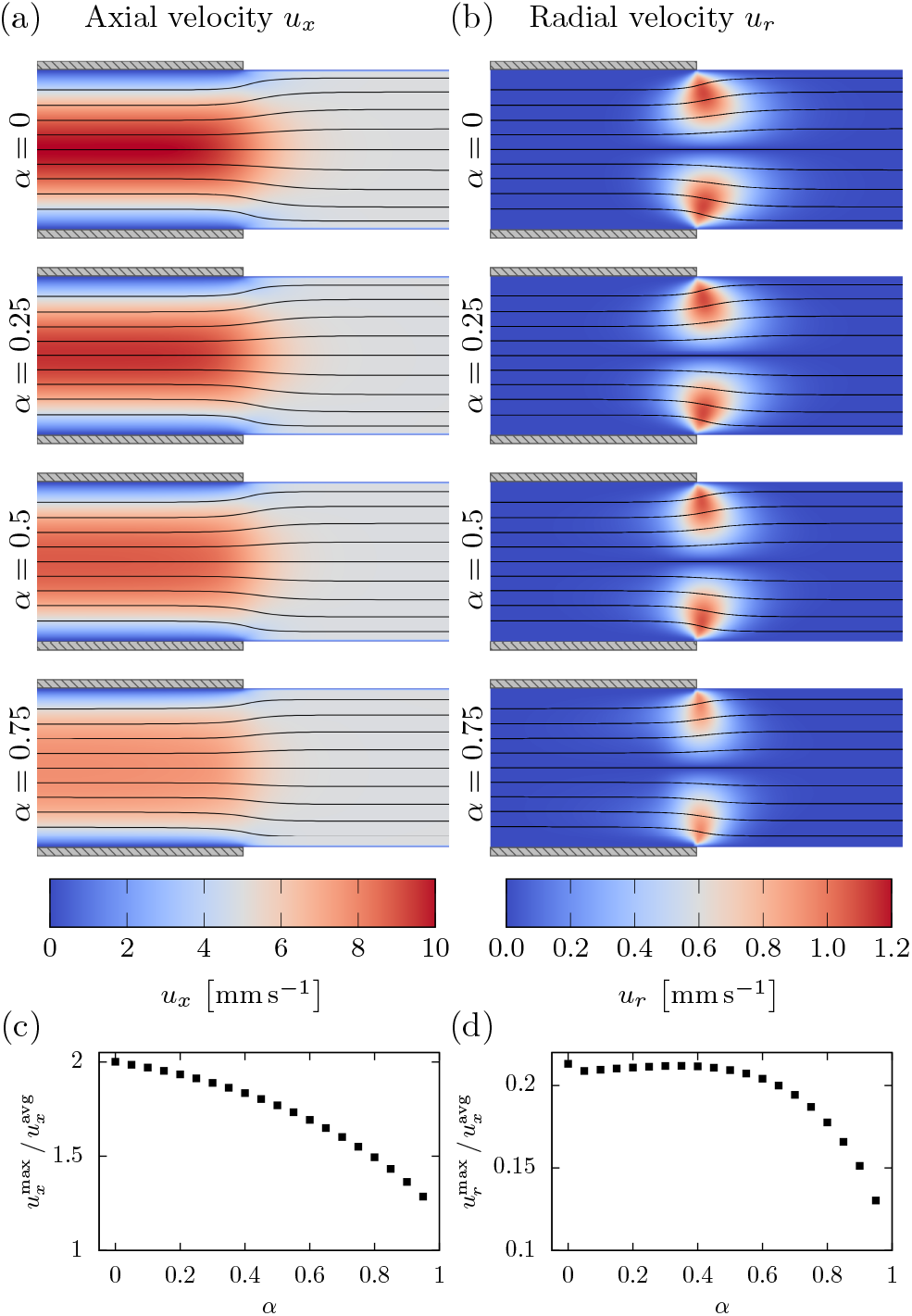
*x*-*y*-slices of the flow at the nozzle exit for increasingly shear thinning fluids: (a) axial velocity component and (b) radial velocity component with streamlines as overlay. (c) The ratio of maximum axial velocity inside the nozzle to the average flow velocity as function of the shear thinning strength *α*. (d) The ratio of maximum radial velocity after the nozzle exit to the average flow velocity as function of *α*.

The second column, figure 7(b), shows the corresponding radial flow components. Due to the radial symmetry, they vanish at the center and increase towards the boundary, showing a drop-like shape with its tip pointing to the position of the nozzle orifice, where the boundary conditions change. The radial flow components decrease with increasing *α*, since the fluid has to be displaced less due to smaller axial velocity difference across the transition. Figure 7(d) quantifies this observation by comparison of the maximum radial flow velocity at the exit with the average axial flow. Combining the axial and radial flows, streamlines are computed in order to visualize the fluid motion in the stationary state. As can be seen in the overlaying lines in figure 7(a) and (b), the streamlines show very similar elongational behavior for all *α* at the nozzle exit due to the simultaneous decrease of the axial and the radial flow component. They are, however, not exactly equal, since the maximum axial and radial velocities scale slightly differently with *α*. Finally, comparing the ratio of axial and radial velocities, we find that the maximum radial flow velocity is always about 10 % of the maximum axial flow velocity, roughly independent of *α*. The fluid stress along the axial direction for different offsets is shown in figure 8 for *α* = 0 and *α* = 0.75. In addition to the total fluid stress, we plot the shear and elongational component separately. To do so, we first decompose the rate of strain tensor into the shear and elongational components

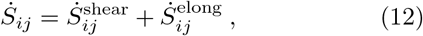

where 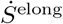 is a diagonal tensor and 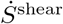 contains only off-diagonal elements. Using this decomposition — further details can be found in section S-3 —, we can define the shear and elongational components of the fluid stress as

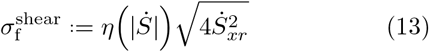

and

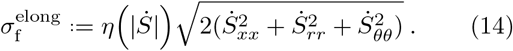

**FIG. 8.**
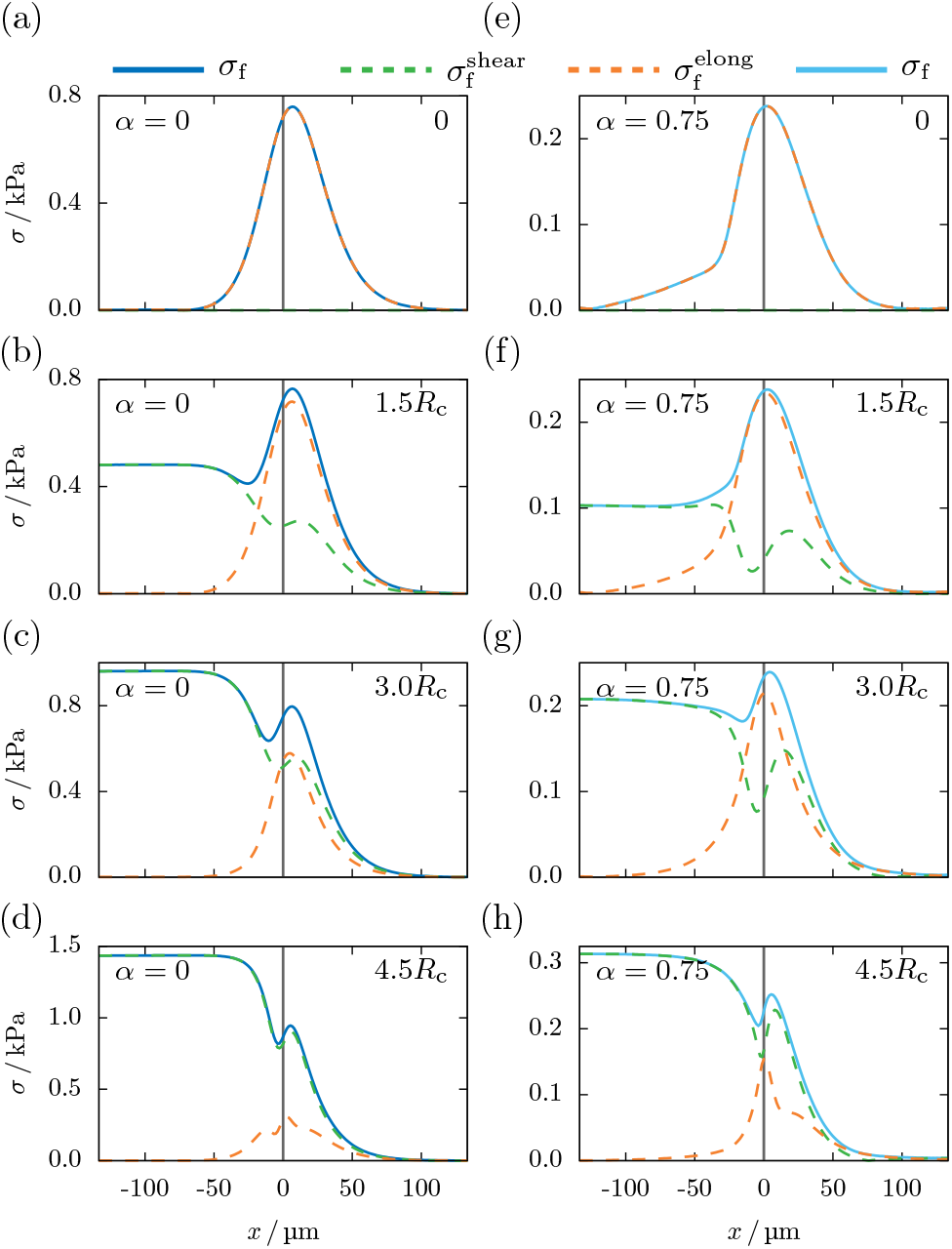
Decomposition of the fluid stress in shear (13) and elongational component (14) at the nozzle exit for the (a-d) Newtonian fluid and (e-h) the highly shear thinning bioink with *α* = 0.75. *x* denotes the axial distance from the nozzle orifice.

Note that 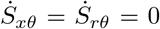, since no azimuthal flow components are present, but that nevertheless 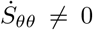 as detailed in section S-3. Thus, the total fluid stress is obtained from (13) and (14) via:

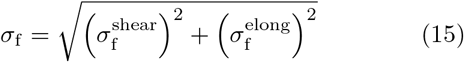

Along the channel center (cf. figure 8(a) and (e)), all shear components of the stress vanish, leaving only the elongational ones, which show a clear peak at the exit. Considering the symmetry, this peak is caused solely by the axial flow deceleration.

With increasing radial offset from the center, as can be seen in figure 8(b-d and f-h) for offsets 1.5*R*_c_, 3.0*R*_c_, and 4.5*R*_c_, the influence of the shear components increases significantly. It can also be seen that the peak of the fluid stress is not only determined by the elongational flow components, but also partly by the shear component 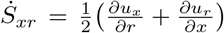. This is further discussed in section S-2 in the SI. The radial offset at which the shear stress inside the nozzle channel exceeds the magnitude of the fluid stress peak depends on the shear thinning strength of the bioink: when comparing figure 8(c) and (g), the stress peak for the Newtonian fluid is already smaller than the fluid stress inside the nozzle channel, while for *α* = 0.75 it is still higher. When selecting shear thinning bioinks in bioprinting, it is thus important to keep in mind that the relative significance of the radial flows at the nozzle exit, both elongational and corresponding shear components, increases when a stronger shear thinning bioink is used.

### D. Cell flowing through the nozzle exit

In this section, we investigate the influence of the flow transition on cells passing the exit of the printer nozzle using our computer simulations and actual micrographs of cells flowing through a real 3D bioprinter nozzle. As discussed in section III C, elongational flow components on a short length scale (≈2*R*) occur at the nozzle exit. These act in different ways on the cell, depending on its radial position when passing the transition:

#### 1. Centered cell

Flowing along the center of the channel, the cell experiences symmetric flow conditions also when passing through the nozzle exit. The deceleration in flow direction leads to an axial compression, while the radial flow stretches the cell in radial direction, leading to an oblate deformation of the cell. As can be seen in the simulation snapshots in figure 9(a) and (b) for the centered flowing cell, its stress uniformly increases inside the whole cell volume during this elongational deformation. After the transition, the cell quickly relaxes towards its spherical equilibrium shape inside the bioink strand.

**FIG. 9.**
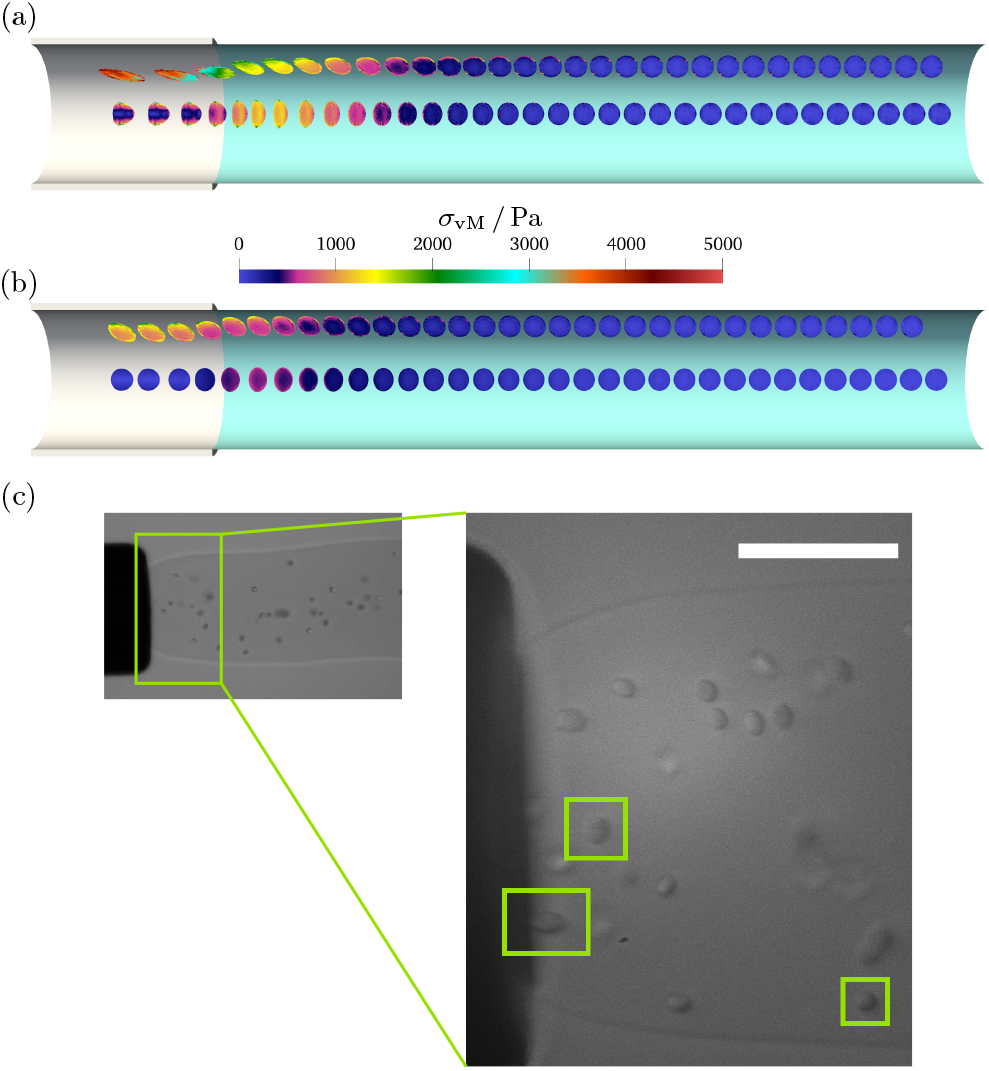
Internal cell stress distribution of cells flowing at different offsets through the nozzle exit in (a) the Newtonian fluid and (b) the shear thinning bioink with *α* = 0.75. Experimental image of cells exiting a 100 µm radius nozzle in 2 % alginate bioink. Left green boxes indicate radially/axially elongated cells flowing in/off-center, respectively. Right green box indicates a cell after relaxation back to equilibrium. Movies of both simulation and experiment can be found in the supplementary material.

Next, we assess the cellular stress resulting from the various flow regimes and ink properties. As can be seen in figure 10(a), an increase in the shear thinning strength of the bioink leads to a decreasing cell stress inside the nozzle channel, as expected from the experimentally observed increased cell survival in more shear thinning bioinks [22, 44]. In contrast to these beneficial effects of shear thinning inside the nozzle, we find that the importance of the elongational stress peak at the nozzle exit notably increases relative to the stress inside the nozzle when *α* is increased: for the Newtonian case (dark blue line figure 10(a)), cell stress increases by approximately 50% from 0.9 kPa to 1.3 kPa during the transition, while for the most shear-thinning bioink (light blue line) it increases six-fold from 0.1 kPa to 0.6 kPa.

**FIG. 10.**
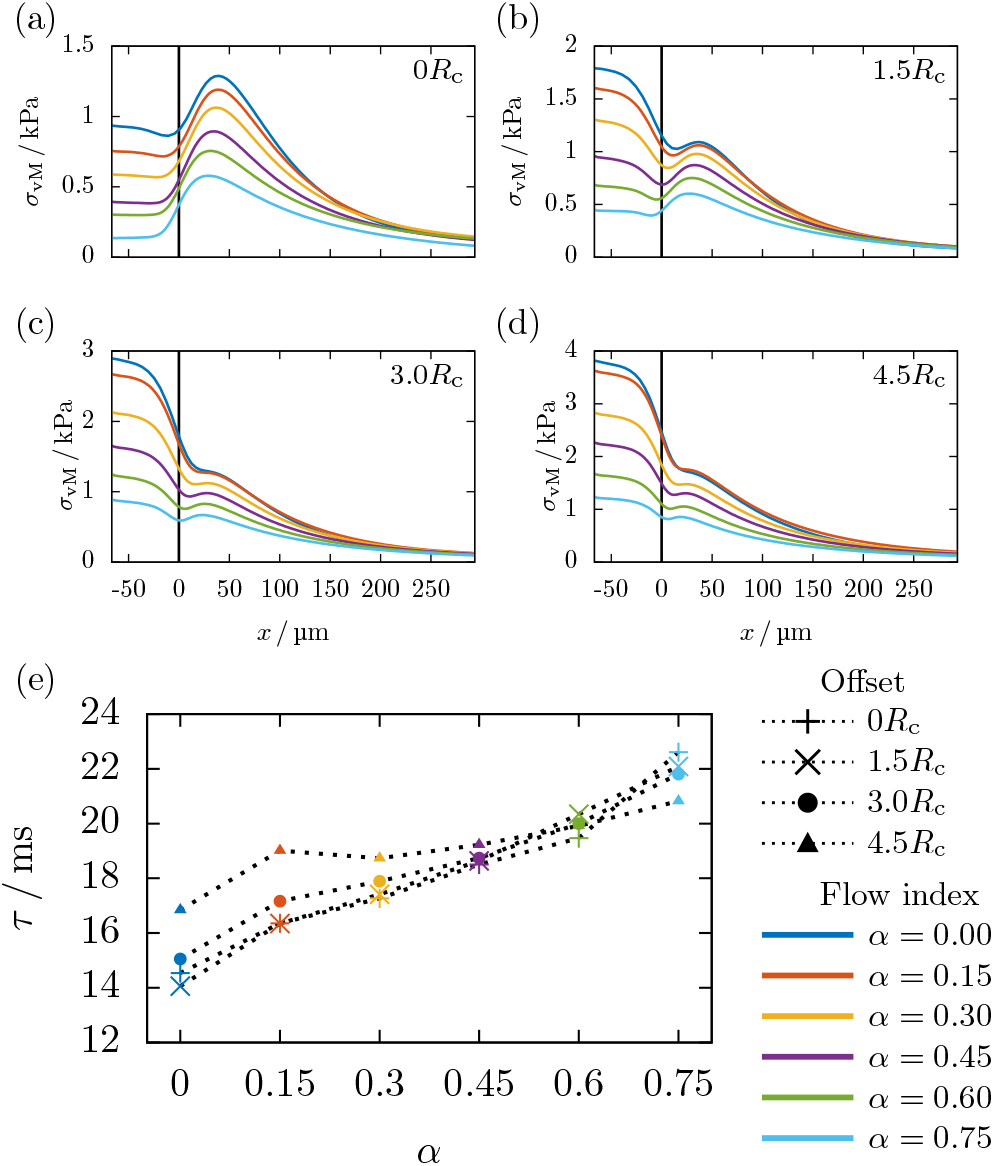
Change of the cell stress when passing through the nozzle exit and flowing in the free bioink strand for increasingly shear thinning bioinks. From (a) to (d), the data is given for the initial cell’s radial offsets 0, 1.5*R*_c_, 3*R*_c_, and 4.5*R*_c_. (e) Relaxation times *τ* of the cell stress when flowing in the free bioink strand as function of *α* and the initial radial cell offset.

Besides cell stresses, an important measure to assess cell damage is cell strain, see e. g. [14]. Due to the symmetry at the channel center, we define an axial strain *α*_1_ := *l*_*x*_*/*(2*R*_c_) and a radial strain *α*_2_ := *l*_*r*_*/*(2*R*_c_), as the maximum elongation of the cell in the considered direction divided by the cell’s reference diameter. As shown in figure 11, the behavior of these cell strains is similar to that of the cell stresses in the paragraph above. Independent of the shear thinning exponent *α*, the axial strain *α*_1_ of the cell’s bullet shape inside the nozzle channel is almost negligible, and only a clear peak in deformation is observed when passing the nozzle exit. The radial strain *α*_2_, on the other hand, already starts with a significant difference from the equilibrium shape. A cell suspended in a highly shear thinning bioink flowing at the nozzle center therefore experiences only the elongational flow right at the nozzle exit, while remaining almost undeformed otherwise.

**FIG. 11.**
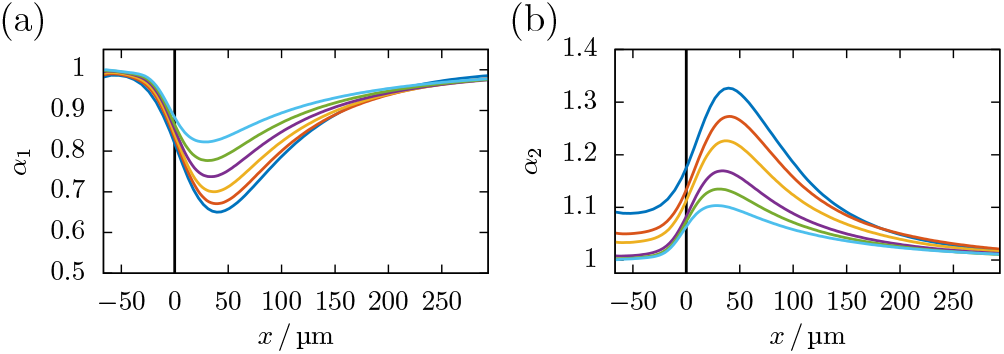
Cell strain for a cell flowing in the center (a) *α*_1_ in *x*-direction and (b) *α*_2_ in *r*-direction at the nozzle exit for the different bioinks. The colors correspond to the flow index as in figure 10.

#### 2. Off-centered cell

We now observe a cell flowing near the nozzle wall Here, the elongational flow at the nozzle exit is combined with shear components inside the nozzle. When passing the transition, the cell is pushed in radial direction leading to a non-ellipsoidal change in shape, before it relaxes towards the equilibrium shape in the bioink strand. An overall decrease of the cell stress when passing through the transition can be observed in the simulation snapshots for the off-centered flowing cell in figure 9(a) and (b). Compared to centered cells in figure 10(a), the importance of elongational relative to shear stress decreases for off-centered cells as shown in figure 10(b)-(d). Indeed, for off-centered cells, the relaxation from the shear-dominated axial flow inside the nozzle channel to the stress-free plug flow in the bioink strand is the most significant effect.

We determine this relaxation time scale *τ* for every simulation by fitting an exponentially decaying function to the cell stress versus time data (see SI figure S-8). Figure 10(e) shows the obtained relaxation times for all cell offsets as function of *α*. We find that the relaxation time increases with increasing shear thinning strength *α* when keeping *η*_0_ constant. This is caused by the larger viscosity of the bioinks with higher *α* for low rates of strain (cf. figure 1), resulting in a higher resistance of the fluid against the cell shape relaxation. Similar to our observations of the fluid stress at the nozzle exit in section III C, we find in figure 10(a to d) that the cell stress peak at the nozzle exit becomes more significant compared to the cell stress inside the nozzle channel when the cell is closer to the center and for stronger shear thinning fluids.

#### 3. Microscopy experiments

To verify our numerical predictions, we image with a high speed camera a bioink strand with cells flowing out of a printing nozzle into a larger reservoir of water. Details of this imaging setup are included in the SI (cf. section S-1). With the objective focused at the tip of the nozzle (inner radius 100 µm), the micrograph in figure 9(c) shows cells suspended in a strand of 2 % alginate bioink during extrusion at a flow rate of 10 µl s^−1^. As can be seen in the marked areas in figure 9(c), cells flowing close to the center exhibit a radially elongated change of shape, while cells flowing near the nozzle wall show an axial elongation. In accordance with our simulations, we observe the cells in the experiment relaxing towards their spherical stress-free shape shortly after the nozzle exit.

### E. Prediction of elongational stress, cell stress and cell strain during bioprinting

The methods employed in sections III B, III C, and III D lead to accurate predictions for important parameters such as cell strain or stress, but require numerical simulations with specialized software. As a practical tool, we develop in the following a simpler yet still accurate method to predict important cell quantities from the printing parameters only.

#### 1. Elongational fluid stress at the nozzle exit

To quantify the importance of elongational effects, we define the average elongational fluid stress 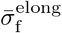 which we obtain by averaging 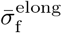 from the simulations along the nozzle axis in an interval of ∓*R* around the peak seen in figure 8(a) and (e). In figure 12 we plot 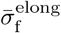 as function of the shear thinning strength of the fluid. As would be expected from the decreasing pressure gradient, the elongational stress monotonously decreases with *α*. In order to obliviate the need for full numerical simulations of the entire flow field in practice, we now show that a good estimate for 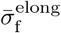 can be obtained by using a much simpler method for flow field computations [41].

**FIG. 12.**
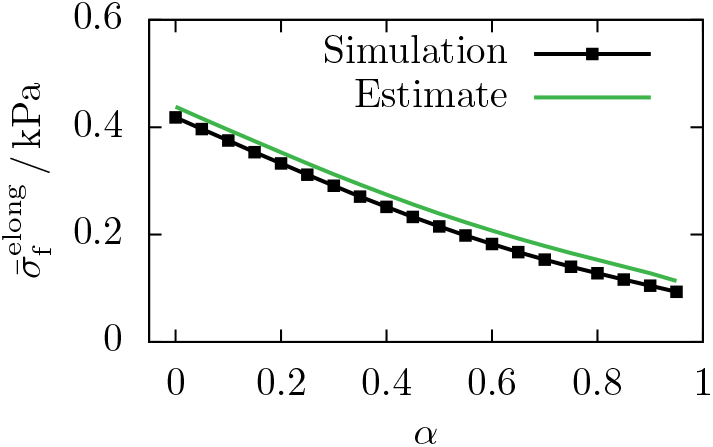
The average elongational stress 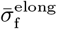 across the nozzle exit from figure 8(a and e) can be estimated from (17).

For this, we assume that the length of the transition is equal to the nozzle diameter 2*R*, as can be verified by comparing with figure 7(a) and (b) and figure 8(a) and (e). Starting from the velocity profile of [41], the change in axial velocity along this length then gives the approximate elongation rate at the nozzle exit:

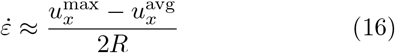

Next, we calculate the stress assuming elongational flow conditions, i. e., 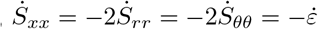, via

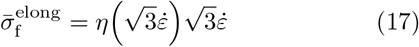

which is derived in the SI (cf. section S-3). This approximated average elongational stress is in very good agreement with the full numerical simulation of the nozzle exit, as shown in figure 12.

We use this approximation to further estimate the elongational cell strain and stress for centered flowing cells at the nozzle exit in the next section.

#### 2. Cell stress and strain for centered cells

We proceed with an estimation of the maximum stress and strain experienced by cells while flowing inside the nozzle as well as during their transition into the free strand at the nozzle exit.

Starting with the latter, we focus on cells flowing at or close to the nozzle center where (as we have shown in figure 8(a), (b), (e), and (f) above) elongational stresses are the most significant fluid stress contribution. The theories of Jeffery and Roscoe [48, 49] contain a solution for the cell strains *α*_1_ and *α*_2_ in a stationary elongational flow (cf. section S-4 B). It reads

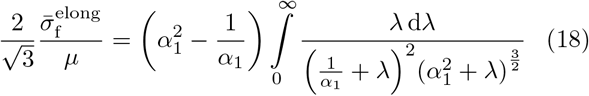

and can be solved numerically for *α*_1_ as function of the elongational fluid stress and the cell’s shear modulus. The other cell strains are 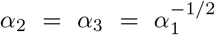 due to symmetry. Using the elongation rate from (17) as input value, we compare the theoretical values with the data obtained from the full numerical simulations in figure 13(a). We note that the theory slightly, but consistently, overestimates cell strains. Indeed, since the elongational flow is experienced by the cell for only a short time span while the theory assumes a stationary elongational flow, this overestimation is to be expected. Interestingly, and in line with what has already been observed in figure 6, Roscoe theory yields surprisingly accurate predictions even for highly shear thinning inks. We again attribute this to the central role of stresses, instead of flow rates, for the cell deformation process in printing nozzles when these are large compared to the radius of the cell. With our approximation consistently over-estimating the simulated results, it can be considered as practical upper limit for predicting cell survival.

**FIG. 13.**
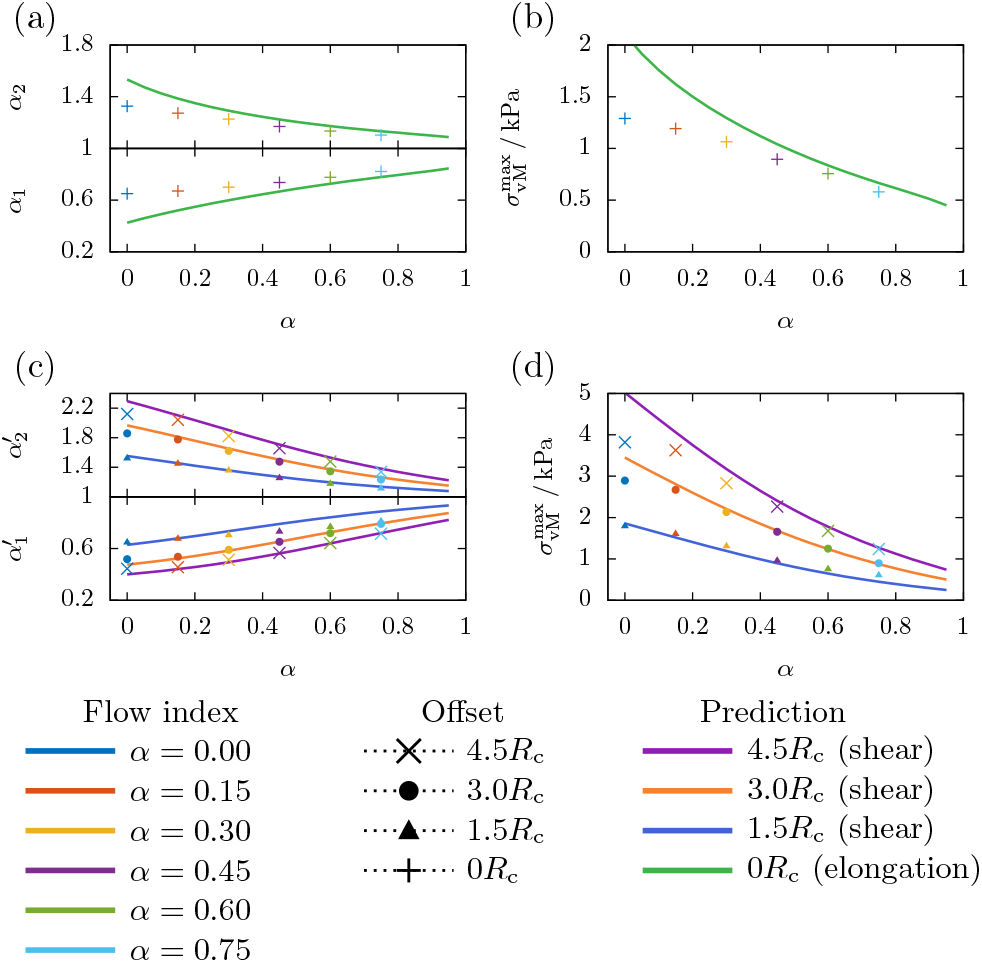
(a) The peak strain and (b) cell stress for centered flowing cells at the transition can be approximated using our estimate of the average elongational fluid stress from (17) and the Jeffery and Roscoe theories for a cell in an elongational flow from section S-4 B. For off-centered cells, our flow calculations inside the nozzle channel and the theory of Roscoe for a cell in shear flow.

As a consequence of the stationarity condition assumed by Roscoe theory, it would predict unrealistically large cell strains in the case of printing velocities higher than the 0.5mm/s used in this work. In reality, however, the flow through the nozzle exit is highly transient and the stationary state is never attained. To assess nevertheless the effect of printing speed, we perform additional simulations for cell flowing centered through the nozzle at 1 cm s^−1^ to 10 cm s^−1^ average extrusion velocity, in order to cover the typical range of 3D bioprinting speeds. Figure 14(a) shows the resulting peak cell strains at the exit from full numerical simulations in comparison to our estimate for 0.5mm/s in figure 13(a). It is apparent that a variation of more than one order of magnitude in flow velocity does hardly affect the cell strains, since the higher velocities significantly decrease the time span during which the high elongational stresses are acting on the cell. Hence, the printing speed does practically not affect the elongational strains occurring during printing.

**FIG. 14.**
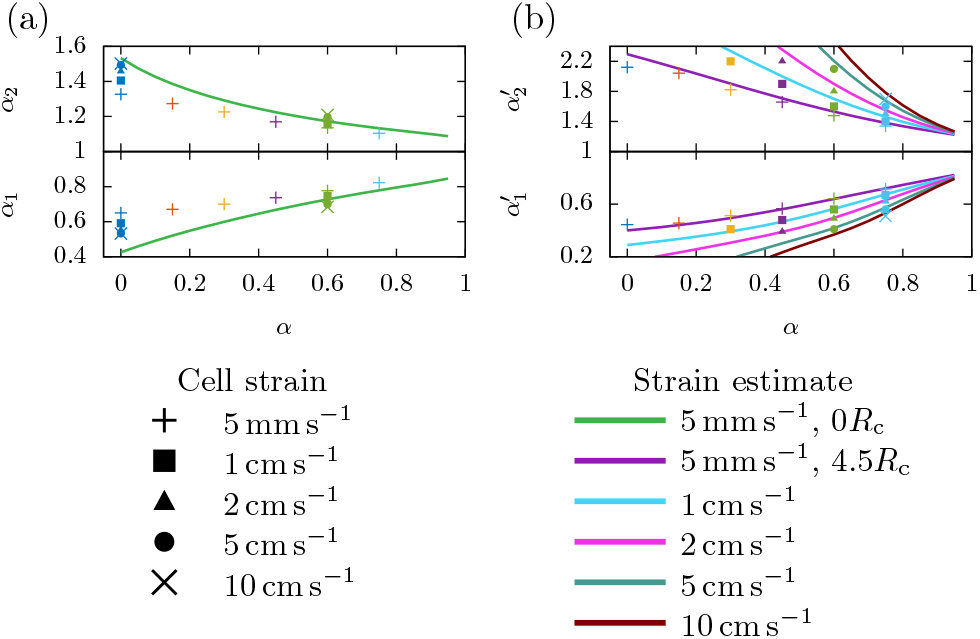
(a) Peak elongational cell strain for centered flowing cells passing the transition for an average extrusion velocity of 1 cm s^−1^, 2 cm s^−1^, 5 cm s^−1^, and 10 cm s^−1^ in comparison to the data of figure 13(a) for 5 mm s^−1^. (b) Maximum cell strain for cells flowing off-centered at 4.5 *R*_c_ for increasing velocities is in accordance with the prediction of the Roscoe theory.

Based on this estimate for cell strain, we proceed to estimate the corresponding cell stress for centered cells. For this, the fluid elongational stress from (17) is fed into the elongational Roscoe theory given by eqs. (18) and (S-52). The result is in good agreement with the full numerical simulations as shown in figure 13(b) for centered cells (green line).

#### 3. Cell stress and strain for off-centered cells

For off-centered cells, we have shown in figure 10(d) that shear components inside the nozzle are an important contribution to the overall cell stress, especially inside less shear thinning bioinks, where they substantially exceed the stress caused by elongational flows at the nozzle exit. We next estimate this overall maximum cell strain and stress.

Due to their almost ellipsoidal shape, we choose as strain measure for the off-centered flowing cells now the ellipsoid’s major and minor semi-axis 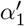 and 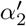, which are obtained through computing the equivalent ellipsoid from the deformed cell’s inertia tensor, as detailed in [30]. Starting from the fluid shear stress obtained from our earlier work [41], we employ the shear part of Roscoe theory in (S-45) and (S-47) and plot the resulting stresses and strains for cells starting at 1.5*R*_c_, 3*R*_c_, and 4.5*R*_c_ in figure 13(c,d). Again, we observe very good agreement with the simulations from section III D(ii).

Upon increasing the average flow velocity by more than one order of magnitude in figure 14(b), we find that cells flowing at maximum radial offset in Newtonian bioinks are not able to attain a stable state while flowing inside the nozzle channel. However, this limitation is solely a result of the large viscosity of the hypothetical Newtonian fluid, and would not affect a real printing process. With increasing shear thinning strength, as shown in figure 14(b), a stable cell shape can be achieved also for high flow velocities of 10 cm s^−1^. The maximum cell strains are accurately predicted by Roscoe theory.

## IV. CONCLUSION

In this work, we investigated the cell stress and strain and the bioink flow behavior during a 3D bioprinting extrusion process using Lattice-Boltzmann numerical simulations together with corresponding qualitative experiments. The two scenarios considered were the flow inside the nozzle channel as well as at the nozzle exit, where the flow transitions into the free bioink strand.

During the first stage of the printing process while cells are flowing inside the printing nozzle, our simulations showed a bullet-like deformation for cells in the center of the channel and an ellipsoidal shape for cells flowing off-center. The latter can be understood on the basis of the classical theory of Roscoe [49] which relates cell stress to the local fluid stress. Interestingly, our simulations demonstrate that these relations hold even in realistic shear thinning bioinks, even though they were originally designed for Newtonian fluids only. The radially inward-directed migration of the cell due to the shear forces was also found to be independent of the shear thinning strength and solely dependent on the printing pressure. We show that, when bioprinting at constant flow rate (or velocity), the shear thinning properties reduce the overall cell stress and strain significantly, while this will not be the case for printing processes performed at constant printing pressure.

In the second stage, cells transition into the free printing strand as they exit the printer nozzle. During this transition, cells are exposed to an elongational flow pattern. While a radial deformation also occurs for cells flowing off-center, we find that the shear deformations dominate in this case. For cells in the channel center, however, this flow causes notable radial stretch of the cells as predicted by our numerical simulations, in qualitative agreement with experimental microscopy images. We show that this effect becomes particularly relevant for cells flowing in highly shear thinning bioinks, as the shear deformation inside the nozzle can virtually be eliminated, while the radial elongation inevitably takes place (figure 10a). In addition, we find that the elongational cell strain is practically independent of the extrusion velocity of the bioink, since the faster velocity balances the high elongational stress by reducing the application time. The relaxation times of the elongated cells even increase with the shear thinning strength, thus prolonging the time that they remain under strain with potentially harmful side effects (figure 10e).

Using our numerical simulation techniques as a starting point together with the velocity profiles derived in our earlier work [41], we finally developed simple estimates for cell stress and/or cell strain for centered as well as off-centered cells.

## Supporting information

Imaging Experiment at the nozzle exit

centered cell in Newtonian bioink

off-centered cell in Newtonian bioink

off-centered cell in shear thinning bioink

centered cell passing the nozzle in Newtonian bioink

centered cell passing the nozzle in shear thinning bioink

off-centered cell passing the nozzle in Newtonian bioink

off-centered cell passing the nozzle in shear thinning bioink

## ACKNOWLEDGMENTS

We thank Nico Schwarm for preparing the cell/alginate sample and help with the imaging experiment. Funded by the Deutsche Forschungsgemeinschaft (DFG, German Research Foundation) — Project number 326998133 — TRR 225 “Biofabrication” (subprojects B07 and A01). We gratefully acknowledge computing time provided by the SuperMUC system of the Leibniz Rechenzentrum, Garching. We further acknowledge support through the computational resources provided by the Bavarian Polymer Institute.

