## Supplementary material for "Predicting cell stress and strain during extrusion bioprinting": Supplemetal Material

#### S-1. BIOPRINTER IMAGING SETUP

MDA-MB-231 breast carcinoma cells were suspended in a 2 % alginate-DMEM solution (sodium alginate PH176, batch nr. 4503283839, JRS Pharma GmbH, Rosenberg, Germany) at a concentration of $10^6\,\frac{\text{cells}}{\text{ml}}$. The cell-alginate suspension was then extruded through a stainless steel needle with an inner diameter of 200 µm and a length of 12.7 mm (Nordson EFD, East Providence, USA) at a constant flow rate of $10\,\mu\text{l}\,\text{s}^{-1}$ using a volume-controlled 3-D printer [1]. The needle was dipped into a transparent plastic cuvette (rotilabo, Roth, Germany) filled with PBS solution. The cells were imaged through a non-infinity corrected 10× 0.25 NA objective (Zeiss, Germany) and a lens-less 150 mm tube (Thorlabs, Germany) using a CMOS camera (acA720-520um, Basler, Germany) at an exposure time of 30 µs and a frame rate of 100 Hz.

FIG. S-1. Decomposition of the shear component of the strain rate tensor for (a) the Newtonian fluid and (b) the bioink with $\alpha = 0.75$. In both cases, the local peak of the rate of strain is due to the radial shear component $\frac{\partial u_r}{\partial x}$, while the axial shear decreases monotonously.

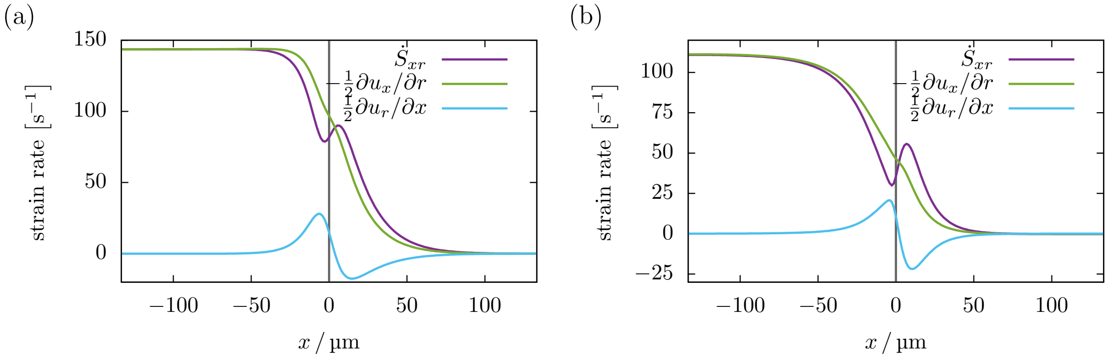


### S-2. FLUID SHEAR STRESS AT THE NOZZLE EXIT

As shown in figure 7 of the manuscript, the total fluid shear stress $\sigma_\mathrm{f}$ along the axial direction always has a local or global peak right after the nozzle exit. When considering the flow close to the channel axis, this peak is clearly a result of the elongational flow at the nozzle exit. Moving closer to the wall, the influence of the elongational components decreases, however, the peak is still present and part of the shear components of the flow. In figure S-1 we decompose the strain rate tensor element $\dot{S}_{xr} = \frac{1}{2}\left(\frac{\partial u_x}{\partial r} + \frac{\partial u_r}{\partial x}\right)$ into its axial and radial shear component. While the axial strain rate $\frac{\partial u_x}{\partial r}$ monotonously decreases along the nozzle exit, the radial strain rate $\frac{\partial u_r}{\partial x}$ increases, changes its sign, and relaxes to zero again due to the localized radial flows at the nozzle exit.

### S-3. FLUID STRESS TENSOR DECOMPOSITION

The components of the rate of strain tensor in a cylindrical coordinate system are given by:

$$\dot{S}_{xx} = \frac{\partial u_x}{\partial x} \tag{S-1}$$

$$\dot{S}_{xr} = \dot{S}_{rx} = \tfrac{1}{2}\left(\frac{\partial u_r}{\partial x} + \frac{\partial u_x}{\partial r}\right) \tag{S-2}$$

$$\dot{S}_{rr} = \frac{\partial u_r}{\partial r} \tag{S-3}$$

$$\dot{S}_{x\theta} = \dot{S}_{\theta x} = \tfrac{1}{2}\left(\frac{\partial u_\theta}{\partial x} + \frac{1}{r}\frac{\partial u_x}{\partial \theta}\right) \tag{S-4}$$

$$\dot{S}_{r\theta} = \dot{S}_{\theta r} = \tfrac{1}{2}\left(\frac{1}{r}\frac{\partial u_r}{\partial \theta} - \frac{u_\theta}{r} + \frac{\partial u_\theta}{\partial r}\right) \tag{S-5}$$

$$\dot{S}_{\theta\theta} = \frac{1}{r}\left(\frac{\partial u_\theta}{\partial \theta} + u_r\right) \tag{S-6}$$

For the present axisymmetric situation, this simplifies to

$$\dot{S}_{ij} = \dot{S}^{\text{shear}}_{ij} + \dot{S}^{\text{elong}}_{ij}\,, \tag{S-7}$$

$$\dot{S}^{\text{shear}} = \begin{pmatrix} 0 & \frac{1}{2}\left(\frac{\partial u_r}{\partial x} + \frac{\partial u_x}{\partial r}\right) & 0 \\ \frac{1}{2}\left(\frac{\partial u_r}{\partial x} + \frac{\partial u_x}{\partial r}\right) & 0 & 0 \\ 0 & 0 & 0 \end{pmatrix}, \tag{S-8}$$

$$\dot{S}^{\text{elong}} = \begin{pmatrix} \frac{\partial u_x}{\partial x} & 0 & 0 \\ 0 & \frac{\partial u_r}{\partial r} & 0 \\ 0 & 0 & \frac{u_r}{r} \end{pmatrix}. \tag{S-9}$$

We can thus compute the scalar shear rate

$$\left|\dot{S}^{\text{shear}}\right| = \sqrt{2\dot{S}^{\text{shear}}_{ij}\dot{S}^{\text{shear}}_{ij}} = \sqrt{4\dot{S}^2_{xr}} \tag{S-10}$$

$$= \left|\frac{\partial u_r}{\partial x} + \frac{\partial u_x}{\partial r}\right| \tag{S-11}$$

and the scalar elongation rate

$$\left|\dot{S}^{\text{elong}}\right| = \sqrt{2\dot{S}^{\text{elong}}_{ij}\dot{S}^{\text{elong}}_{ij}} \tag{S-12}$$

$$= \sqrt{2(\dot{S}^2_{xx} + \dot{S}^2_{rr} + \dot{S}^2_{\theta\theta})} \tag{S-13}$$

$$= \sqrt{2\left(\frac{\partial u_x}{\partial x}\right)^2 + 2\left(\frac{\partial u_r}{\partial r}\right)^2 + 2\left(\frac{u_r}{r}\right)^2}\,, \tag{S-14}$$

and the rate of strain

$$\left|\dot{S}\right| = \sqrt{\left|\dot{S}^{\text{shear}}\right|^2 + \left|\dot{S}^{\text{elong}}\right|^2}\,. \tag{S-15}$$

Using the shear and elongation rates, we define the fluid's scalar shear and elongational stress via

$$\sigma_{\text{f}}^{\text{shear}} := \eta\left(|\dot{S}|\right)\left|\dot{S}^{\text{shear}}\right| \tag{S-16}$$

and

$$\sigma_{\text{f}}^{\text{elong}} := \eta\left(|\dot{S}|\right)\left|\dot{S}^{\text{elong}}\right|\,. \tag{S-17}$$

The total fluid stress is thus obtained as:

$$\sigma_{\text{f}} = \eta\left(|\dot{S}|\right)|\dot{S}| = \sqrt{\left(\sigma_{\text{f}}^{\text{shear}}\right)^2 + \left(\sigma_{\text{f}}^{\text{elong}}\right)^2} \tag{S-18}$$

When assuming a perfect elongational flow, i. e., $u_x = -\dot{\varepsilon}x$, $u_r = \frac{1}{2}\dot{\varepsilon}r$, and $u_\theta = 0$, the fluids rate of strain is given via

$$|\dot{S}| = \left|\dot{S}_{ij}^{\text{elong}}\right| = \sqrt{2\dot{\varepsilon}^2 + 2\left(\tfrac{1}{2}\dot{\varepsilon}\right)^2 + 2\left(\tfrac{1}{2}\dot{\varepsilon}\right)^2} = \sqrt{3}\dot{\varepsilon} \tag{S-19}$$

and thus follows the fluid stress as:

$$\sigma_{\text{f}} = \sigma_{\text{f}}^{\text{elong}} = \eta\left(\sqrt{3}\dot{\varepsilon}\right)\sqrt{3}\dot{\varepsilon} \tag{S-20}$$

### S-4. JEFFERY AND ROSCOE THEORY

An analytical theory describing the deformation and stresses of a cell embedded in a linear flow was proposed by Roscoe [2], based on the work of Jeffery [3]. For convenience, we briefly summarize their theoretical approach and the application of the theory for cells in a shear and an elongational flow scenario in this section.

Jeffery [3] originally solved the problem of the motion of a rigid ellipsoidal particle in a linear flow, i. e., the undisturbed fluid velocity can be written as (using the notation of Roscoe [2])

$$v'_i = e'^{(1)}_{ij} x_j - \zeta'_{ij} x_j \,, \tag{S-21}$$

where the fluid's rate of strain and vorticity are defined by

$$e'^{(1)}_{ij} = \frac{1}{2}\left(\frac{\partial v'_i}{\partial x_j} + \frac{\partial v'_j}{\partial x_i}\right) \quad \text{and} \quad \zeta'_{ij} = \frac{1}{2}\left(\frac{\partial v'_i}{\partial x_j} - \frac{\partial v'_j}{\partial x_i}\right) . \tag{S-22}$$

Jeffery [3] derived the fluid stress acting on the surface of a rigid ellipsoidal particle as

$$p'_{ij} = -p_\mathrm{h}\delta_{ij} + \eta_0 A_{ij} \,, \tag{S-23}$$

with an arbitrary hydrostatic pressure $p_\mathrm{h}$ and a deviatoric tensor $A_{ij}$. Roscoe [2] notes that the deviatoric stress can further be divided into two parts,

$$p'_{ij} = -p_\mathrm{h}\delta_{ij} + 2\eta_0 e'^{(1)}_{ij} + \eta_0\left(A_{ij} - 2e'^{(1)}_{ij}\right) , \tag{S-24}$$

one due to the undisturbed flow from (S-21) and one due to the disturbance of the flow induced by the particle presence. The components $A_{ij}$ in a coordinate system coinciding with the ellipsoid axes can be calculated via (remaining components by cyclic change of indices):

$$A_{11} = \frac{4}{3}\frac{2g''_1 e'^{(1)}_{11} - g''_2 e'^{(1)}_{22} - g''_3 e'^{(1)}_{33}}{g''_2 g''_3 + g''_3 g''_1 + g''_1 g''_2} \tag{S-25}$$

$$A_{12} = \frac{g_1 e'^{(1)}_{12} - \alpha_2^2 g'_3 \zeta'_{12}}{2g'_3(\alpha_1^2 g_1 + \alpha 2^2 g_2)} \tag{S-26}$$

$g_i$, $g_i'$, and $g_i''$ are integrals of the type

$$g_1 = \int_0^\infty \frac{\mathrm{d}\lambda}{(\alpha_1^2+\lambda)\Delta} \tag{S-27}$$

$$g_1' = \int_0^\infty \frac{\mathrm{d}\lambda}{(\alpha_2^2+\lambda)(\alpha_3^2+\lambda)\Delta} = \frac{g_3-g_2}{\alpha_2^2-\alpha_3^2} \tag{S-28}$$

$$g_1'' = \int_0^\infty \frac{\lambda\,\mathrm{d}\lambda}{(\alpha_2^2+\lambda)(\alpha_3^2+\lambda)\Delta} = \frac{\alpha_2^2 g_2-\alpha_3^2 g_3}{\alpha_2^2-\alpha_3^2}\,, \tag{S-29}$$

where $\Delta = \sqrt{(\alpha_1^2+\lambda)(\alpha_2^2+\lambda)(\alpha_3^2+\lambda)}$.

Equation (S-23) can directly be employed to compute the stresses acting on a rigid ellipsoid suspended in the undisturbed flow given by (S-21), e. g., a cell inside an elongational flow, as detailed in section S-4 B. Roscoe [2] extended the theory of Jeffery [3] to compute the stresses acting on a non-rigid ellipsoid with moving boundaries, i. e., tank-treading motion. The ellipsoid's surface motion is assumed to be linear — similar to (S-21) — given by

$$v_i = \bar{e}_{ij}^{(1)} x_j - \bar{\zeta}_{ij} x_j\,, \tag{S-30}$$

with $\bar{e}_{ij}^{(1)}$ and $\bar{\zeta}_{ij}$ denoting respectively the average rate of strain and vorticity inside the particle, which are always equal to their values at the particle surface [2]. The velocity disturbance $\Delta v_i' = v_i - v_i'$ at the particle surface induced by the surface motion of the non-rigid particle is equal to the velocity disturbance of a rigid particle in an undisturbed flow given by

$$v_i'' = v_i' - v_i\,. \tag{S-31}$$

Therefore, (S-24) can be employed to compute the fluid stresses for a non-rigid particle with a moving boundary by simply computing the stress contribution due to the disturbance using the equivalent undisturbed flow (S-31), while keeping the contribution due to the actual undisturbed flow (S-21). Thus:

$$p_{ij}' = -p_\mathrm{h}\delta_{ij} + 2\eta_0 e_{ij}'^{(1)} + \eta_0\Big(A_{ij}' - 2\Big(e_{ij}'^{(1)} - \bar{e}_{ij}^{(1)}\Big)\Big) \tag{S-32}$$

$$= -p_\mathrm{h}\delta_{ij} + \eta_0\Big(A_{ij}' + 2\bar{e}_{ij}^{(1)}\Big) \tag{S-33}$$

Here, the tensor $A_{ij}'$ is computed for an undisturbed flow of the form given in (S-31) instead of (S-21).

#### A. Cell stress and strain in shear flow

Roscoe [2] applies (S-32) to compute the motion of a tank-treading ellipsoidal particle in a linear shear flow. The coordinates of a material point of the particle starting at position $(\tilde{x}_1, \tilde{x}_2, \tilde{x}_3)$ following an elliptical trajectory are given by

$$x_1 = \alpha_1(\tilde{x}_1 \cos(\nu t) - \tilde{x}_2 \sin(\nu t)) \tag{S-34}$$

$$x_2 = \alpha_2(\tilde{x}_1 \sin(\nu t) + \tilde{x}_2 \cos(\nu t)) \tag{S-35}$$

$$x_3 = \alpha_3 \tilde{x}_3 \,, \tag{S-36}$$

where $x_1$, $x_2$, and $x_3$, align with the ellipsoid's semi-axes, thus yielding the surface velocity:

$$v_1 = -\frac{\alpha_1}{\alpha_2} \nu x_2 \tag{S-37}$$

$$v_2 = \frac{\alpha_2}{\alpha_1} \nu x_1 \tag{S-38}$$

$$v_3 = 0 \tag{S-39}$$

The surface velocity defines the rate of strain and vorticity from (S-30). A linear shear flow — commonly described in the global coordinate system as $v_1' = \kappa x_2$, $v_2' = v_3' = 0$ with a shear rate $\kappa$ — written in terms of a coordinate system aligned with the ellipsoid's semi-axes through rotation by an angle $\theta$ is given by

$$v_1' = \kappa\big(x_1 \sin\theta \cos\theta + x_2 \cos^2(\theta)\big) \tag{S-40}$$

$$v_2' = -\kappa\big(x_1 \sin^2(\theta) + x_2 \sin\theta \cos\theta\big) \tag{S-41}$$

$$v_3' = 0 \,. \tag{S-42}$$

From that, the undisturbed fluid's rate of strain and vorticity from (S-21), and, together with (S-30), the fluid stress at the particle surface from (S-32) can be computed.

In a stationary state, the fluid stress must be balanced by the cell stress at the particle surface. As mentioned in section III-A of the manuscript, the cell stress consists of an elastic and a viscous part. For the triaxial ellipsoidal deformation described in (S-34), the elastic stress at the particle surface can be computed from (9) assuming an incompressible cell ($J = 1$). With the corresponding deformation gradient tensor given by $F_{ij} = \alpha_i \delta_{ij}$, the non-zero diagonal elements of the Cauchy stress are then found as

$$\sigma_{11} = \frac{\mu}{3}\left(2\alpha_1^2 - \alpha_2^2 - \alpha_3^2\right) \,, \tag{S-43}$$

with similar expressions for $\sigma_{22}$ and $\sigma_{33}$ obtained by cyclic change of indices. The obtained system of two equations of the stress balance is solved by considering only the differences of the principal stresses, which eliminates the hydrostatic pressure:

$$p'_{11} - p'_{22} = \sigma_{11} - \sigma_{22} \tag{S-44}$$

$$\Leftrightarrow 2\eta_0\kappa\sin(2\theta)\frac{g''_1 + g''_2}{g''_2g''_3 + g''_3g''_1 + g''_1g''_2} = \mu\left(\alpha_1^2 - \alpha_2^2\right) \tag{S-45}$$

$$p'_{11} + p'_{22} - 2p'_{33} = \sigma_{11} + \sigma_{22} - 2\sigma_{33} \tag{S-46}$$

$$\Leftrightarrow 2\eta_0\kappa\sin(2\theta)\frac{g''_1 - g''_2}{g''_2g''_3 + g''_3g''_1 + g''_1g''_2} = \mu\left(\alpha_1^2 + \alpha_2^2 - \frac{2}{\alpha_1^2\alpha_2^2}\right) \tag{S-47}$$

Note that $\alpha_3 = \frac{1}{\alpha_1\alpha_2}$ due to the assumed incompressibility. The viscous contribution of the cell stress can be computed directly from its internal fluid motion (S-37)–(S-39) as:

$$\sigma_{12} = \sigma_{21} = 2\eta_1 e'^{(1)}_{12} = -\eta_1\nu\frac{\alpha_1^2 - \alpha_2^2}{\alpha_1\alpha_2} \tag{S-48}$$

Through numerical solution of these equations one obtains the cell stresses and strains as well as the tank-treading frequency as function of the undisturbed fluid's shear rate $\kappa$ (cf. Roscoe[2, eq.(80),(41)]).

To compare it with the Roscoe theory, we numerically assess the viscous shear stress inside the cell from our simulations by extracting the Lattice-Boltzmann strain rate tensor field [4, 5] inside the cell using our method from [6] and averaging over the cell volume. In figure S-2 we show how the resulting viscous component of the cell stress as function of the cell's shear modulus in a linear shear flow of constant strain rate $|\dot{S}| = 100\,\mathrm{s}^{-1}$. For low $\mu$ — i. e., for soft cells — the shear stress inside the cells asymptotically approaches the viscous shear stress of the surrounding undisturbed fluid, as it is increasingly stretched and hence more aligned with the flow. Very stiff cells, on the other hand, will remain their undeformed spherical shape. It can be seen that the transition between these two limits in large parts happens in the stiffness range of biological cells at around 100 Pa to 10 kPa.

FIG. S-2. Viscous shear stress $\sigma_{12}$ inside the cell as function of the cell stiffness in a shear flow with $|\dot{S}| = 100\,\mathrm{s}^{-1}$. The straight line indicates the viscous fluid stress of the surrounding undisturbed flow field.

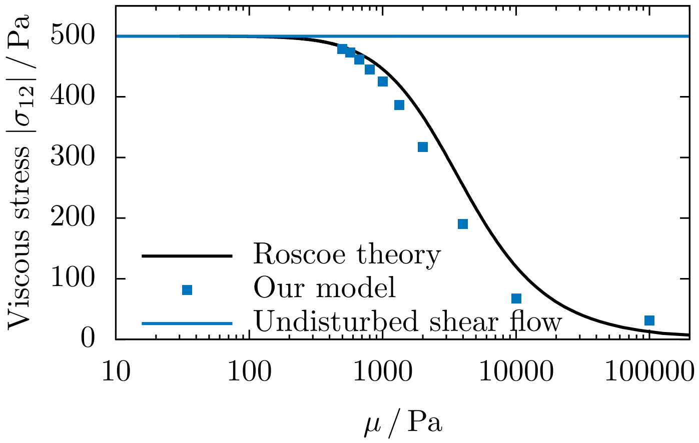


#### B. Cell stress and strain in elongational flow

Roscoe [2] applies (S-23) to compute the steady state deformation and stresses of an ellipsoid in an elongational flow, where the undisturbed velocity is given by

$$v'_1 = \xi x_1 \tag{S-49}$$

$$v'_2 = -\frac{1}{2}\xi x_2 \tag{S-50}$$

$$v'_3 = -\frac{1}{2}\xi x_3\,, \tag{S-51}$$

with the elongational rate $\xi$. From this, the rate of strain and vorticity in (S-21) and the deviatoric tensor $A_{ij}$ are calculated. Since the ellipsoid's boundary has no motion, $v_i = 0$. Due to the symmetry of the flow and the incompressibility, $\alpha_2 = \alpha_3 = \alpha_1^{-1/2}$ applies to the ellipsoid. The fluid's normal stress differences from (S-23) are then set to balance the surface stresses of the triaxially elongated particle:

$$p'_{11} - p'_{22} = \sigma_{11} - \sigma_{22} \tag{S-52}$$

$$2\eta_0 \frac{\xi}{g''_2} = \mu\left(\alpha_1^2 - \frac{1}{\alpha_1}\right) \tag{S-53}$$

The numerical solution of this equation yields the cell stresses and strains as function of the undisturbed fluid's elongational rate $\xi$. We note that due to the stationarity condition a stable solution can not be found for very high elongational rates.

### S-5. APPLICABILITY OF ROSCOE THEORY FOR SHEAR THINNING BIOINKS

In figure 5(c and d) of the manuscript we show the cell stress as function of the fluid stress for different cell starting positions in the channel in a Newtonian bioink and for the maximum radial offset for increasing shear thinning strength. Figure S-3 shows additional data curves for all investigated bioinks, i. e., data similar to figure 5(c) for different $\alpha$. In Addition to the fluid stress on the lower $x$-axis, the upper $x$-axis gives the radial position of the cell in units of the cell radius.

As mentioned in section III A of the manuscript, the key property determining cell motion is the shear stress. To underline this, we plot in figure S-4 the cell stress data from figure 5(d), but with respect to the rate of strain instead of the shear stress. Due to the similar velocities, the range of the $|\dot{S}|$-axis is similar for all flow indices. Therefore — instead of collapsing onto a master curve as in figure 5(d) — the curves fan out, suggesting a weaker dependency of the cell stress on the shear rate for increasingly shear thinning bioinks. This, however, is slightly misleading, since it neglects the change in viscosity of the surrounding liquid.

In section III E we find that the influence of higher extrusion velocities on the elongational cell strain is almost negligible. However, this does obviously not apply to the shear conditions inside the nozzle channel, as a higher pressure gradient is necessary to produce larger flow velocities. In figure S-5 below, we show data similar to that of figure 5(c and d), for a cell starting at the largest radial offset in a bioink with $\alpha = 0.6$ for average extrusion velocities of $1\,\mathrm{cm\,s^{-1}}$, $2\,\mathrm{cm\,s^{-1}}$, and $5\,\mathrm{cm\,s^{-1}}$, demonstrating the validity of the Roscoe theory also for higher velocities.

FIG. S-3. The cell stress inside the nozzle channel as function of the local shear stress for all used bioinks, as in figure 5(c). The upper $x$-axis gives the radial position of the cell in units of the cell radius.

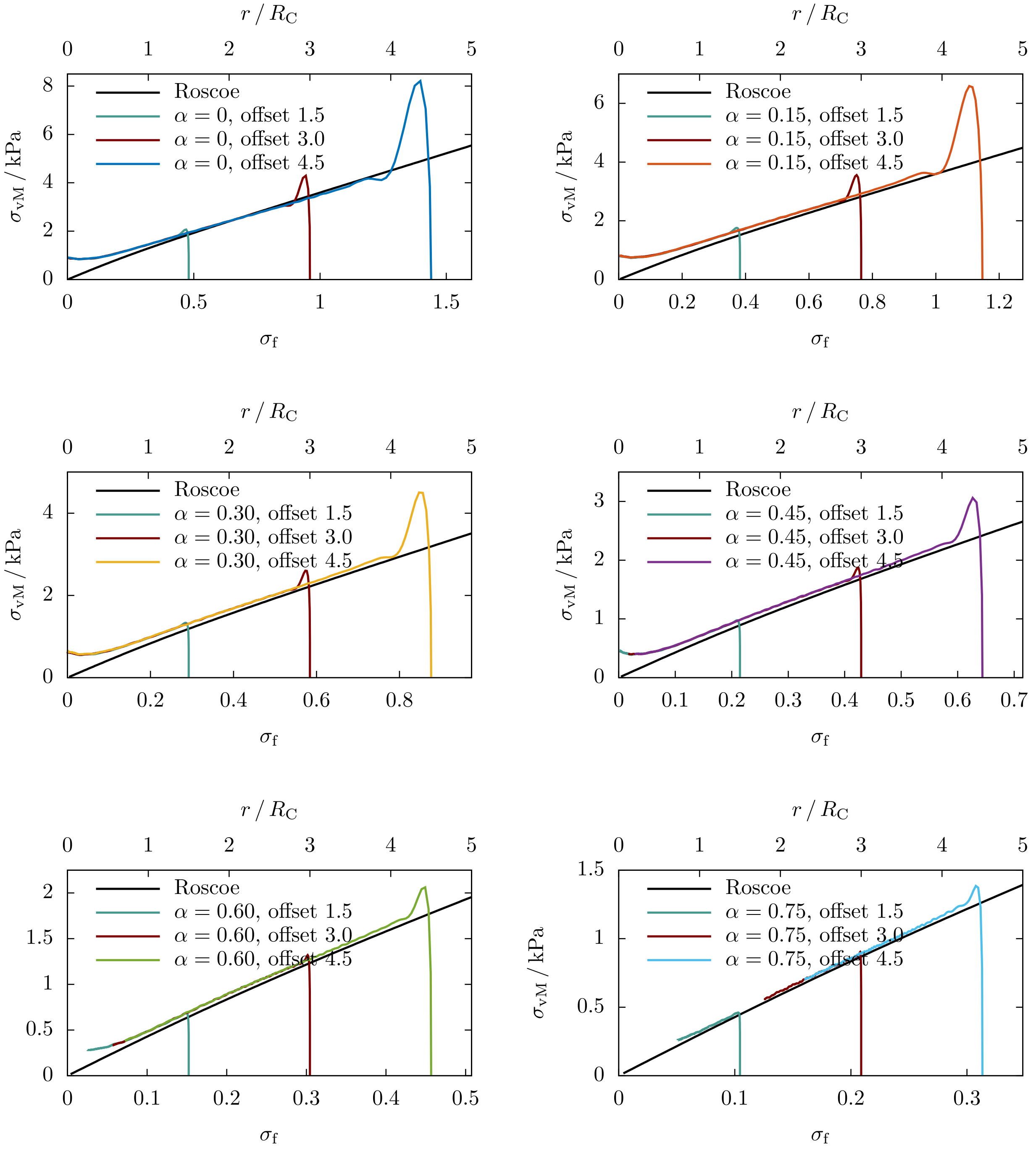

FIG. S-4. Data from figure 5(d), but plotted versus the local rate of strain $|\dot{S}|$ of the fluid. Due to the constant average velocity, the shear rates experienced by the cells are of similar magnitude.

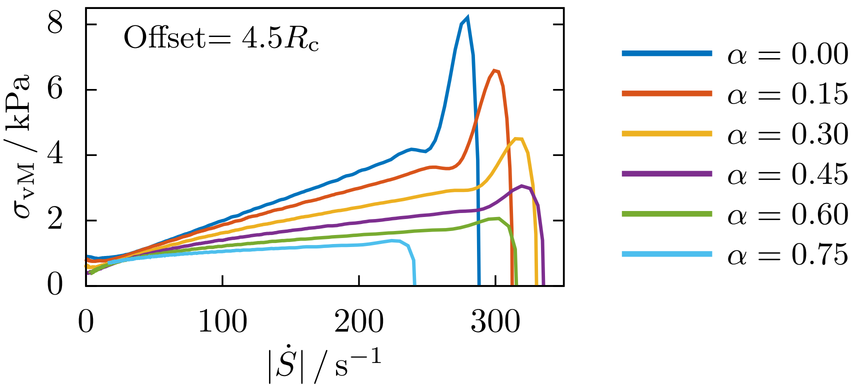


FIG. S-5. The cell stress as function of the fluid stress (similar to figure 5(d) for a cell starting at the largest radial offset in a bioink with $\alpha = 0.6$ for average extrusion velocities of $1\,\mathrm{cm\,s^{-1}}$, $2\,\mathrm{cm\,s^{-1}}$, and $5\,\mathrm{cm\,s^{-1}}$, demonstrating the validity of the Roscoe theory also for higher velocities.)

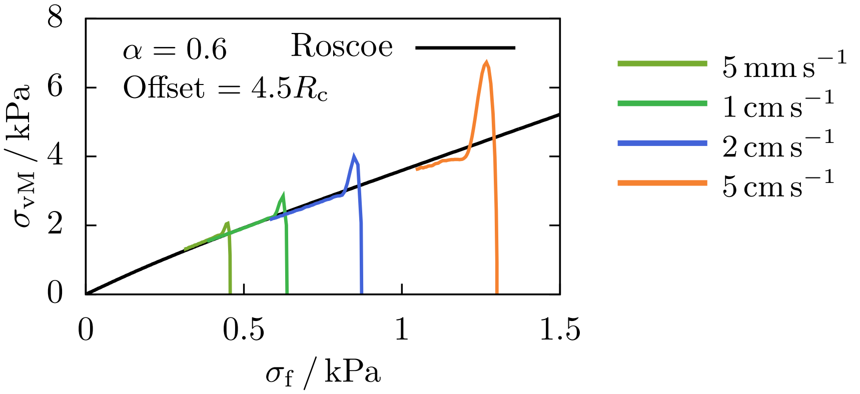

### S-6. RIGID SPHERE IN FLOW

To compute the additional stress caused by a rigid sphere in pure shear flow, we start from the strain rate tensor (S-22) of the undisturbed flow. As usual, for a shear rate $\kappa$, this is given by

$$\underline{e}'^{(1)} = \frac{1}{2}\begin{pmatrix} 0 & \kappa & 0 \\ \kappa & 0 & 0 \\ 0 & 0 & 0 \end{pmatrix} \tag{S-54}$$

The sphere is neutrally buouyant as well as force- and torque-free. We then compute the so-called stresslet (see, e. g., [7, eq. (2.32)]) which embodies the additional stress in the fluid due to the presence of the sphere

$$\begin{aligned} \underline{S} &= \frac{20}{3}\pi\eta_0 R_c^3 \underline{e}'^{(1)} \\ &= \frac{20}{6}\pi\eta_0 R_c^3 \kappa \begin{pmatrix} 0 & 1 & 0 \\ 1 & 0 & 0 \\ 0 & 0 & 0 \end{pmatrix} \end{aligned} \tag{S-55}$$

This quantity is normalized by the shear stress of the undisturbed fluid integrated over the sphere volume

$$S_f = \frac{4}{3}\pi R_c^3 \eta_0 \kappa \tag{S-56}$$

thus leading to the dimensionless stresslet

$$\begin{aligned} \underline{S}^* &= \frac{\underline{S}}{S_f} \\ &= \frac{5}{2}\begin{pmatrix} 0 & 1 & 0 \\ 1 & 0 & 0 \\ 0 & 0 & 0 \end{pmatrix}. \end{aligned} \tag{S-57}$$

(S-57) is given in the laboratory system. To express it in the body-fixed coordinate system of the cell, we require a rotation by 45° (cf. left inset in figure 4) given by the matrix

$$\underline{M}_{45} = \frac{\sqrt{2}}{2}\begin{pmatrix} 1 & -1 & 0 \\ 1 & 1 & 0 \\ 0 & 0 & 1 \end{pmatrix}. \tag{S-58}$$

The final result is

$$\underline{S}^*_{\mathrm{rot}} = \underline{M}_{45}\ \underline{S}^*\ \underline{M}^{\mathrm{T}}_{45}$$
$$= \frac{5}{2}\begin{pmatrix} -1 & 0 & 0 \\ 0 & 1 & 0 \\ 0 & 0 & 0 \end{pmatrix} \tag{S-59}$$

thus furnishing an explanation for the cell stress at low flow rates in figure 4.

### S-7. UNIAXIAL STRETCHING OF AN ELASTIC BEAM

In the limit of high Capillary numbers, the elastic components of the cell stress tensor approximately develop according to the ratio $\sigma_{11} : \sigma_{22} : \sigma_{33} = 2 : -1 : -1$. This ratio is equivalent to what would be expected from the uniaxial extension of an elastic beam, as briefly outlined in the following. The uniaxial stretching with a factor $\alpha_1 = a$ in $x_1$-direction of an isotropic, incompressible material results in $\alpha_2 = \alpha_3 = \frac{1}{\sqrt{a}}$ for the remaining principal stretches. The left Cauchy-Green deformation tensor is hence given by $B = \mathrm{diag}\big(a^2, \frac{1}{a}, \frac{1}{a}\big)$, which can be inserted into (S-43) of the manuscript in order to obtain the stress components as:

$$\sigma_{11} = \frac{2}{3}\mu\left(a^2 - \frac{1}{a}\right) \tag{S-60}$$

$$\sigma_{22} = \sigma_{33} = -\frac{1}{3}\mu\left(a^2 - \frac{1}{a}\right) \tag{S-61}$$

### S-8. STRESS RELAXATION INSIDE THE BIOINK STRAND

In figure S-6 we show the individual fits of the cell stress relaxation times $\tau$ from figure 9(e) of the manuscript. As a fit function we use an exponential decay of the form

$$\sigma_{\mathrm{vM}}(t) = \sigma_{\mathrm{vM}}^{\mathrm{arb.offset}} + \sigma_{\mathrm{vM}}^{(0)} \exp\left(-\frac{t-t_0}{\tau}\right) , \tag{S-62}$$

where $\sigma_{\mathrm{vM}}^{(0)}$ denotes the cell stress at $t_0$ (indicated by the gray area), and $\sigma_{\mathrm{vM}}^{\mathrm{arb.offset}}$ is an arbitrary small offset. At $t_0$, the cell passes the transition (i. e.$x = 0$).

When the cell relaxes in a quiescent fluid, i. e., when we disable any imposed flow or external pressure, the relaxation times of the cell decrease slightly. This is shown in figure S-7, where we compare the relaxation times of cells inside quiescent fluid to those of cells passing through the nozzle exit into the bioink strand (cf.figure 9(e))

FIG. S-6. Relaxation time fit of the cell stress in the bioink strand using an exponentially decaying function. The fit excludes they gray shaded area.

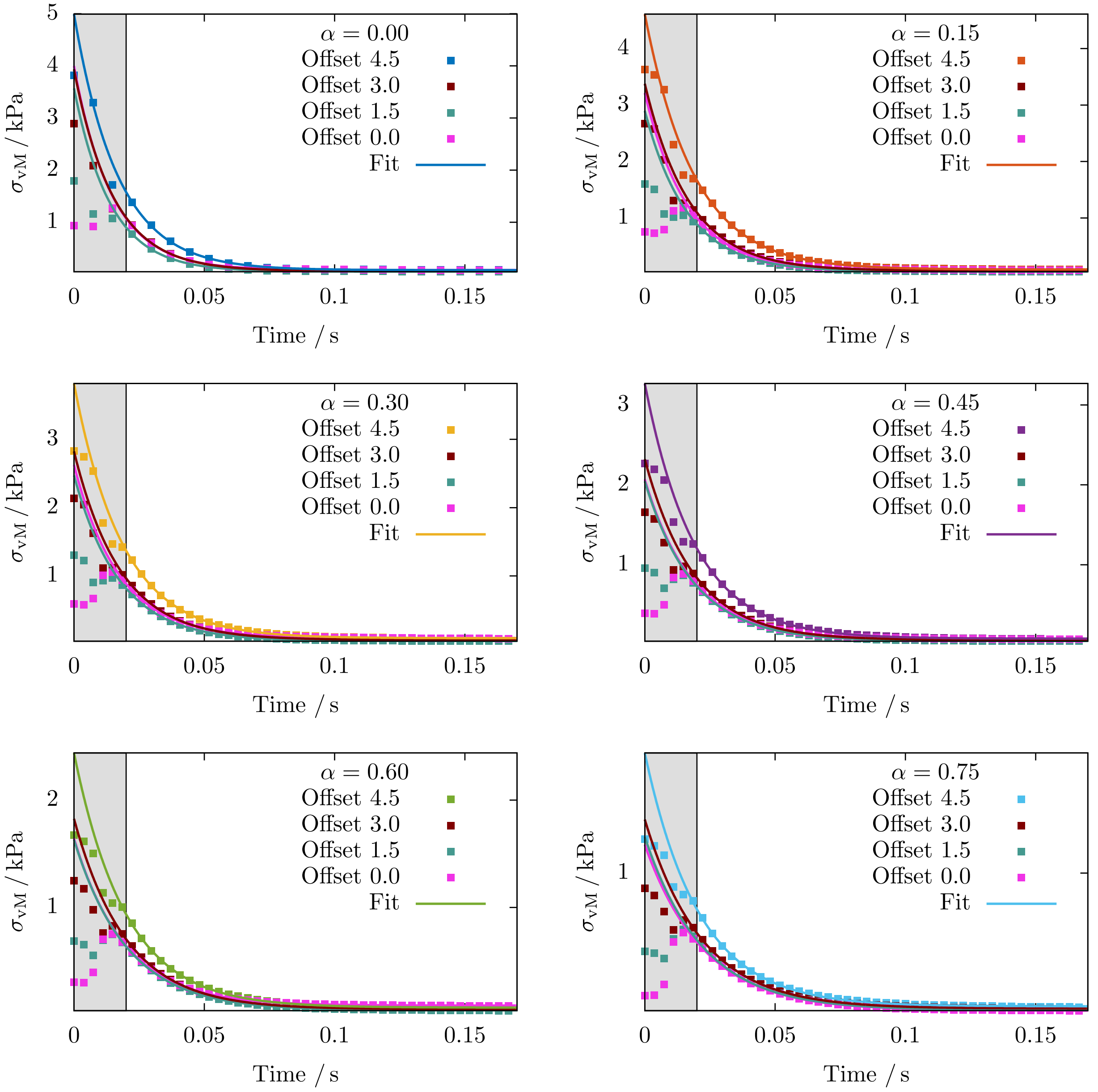

FIG. S-7. Relaxation times of cells suspended in quiescent liquid (solid lines), compared to the data in figure 9(e) (dotted lines).

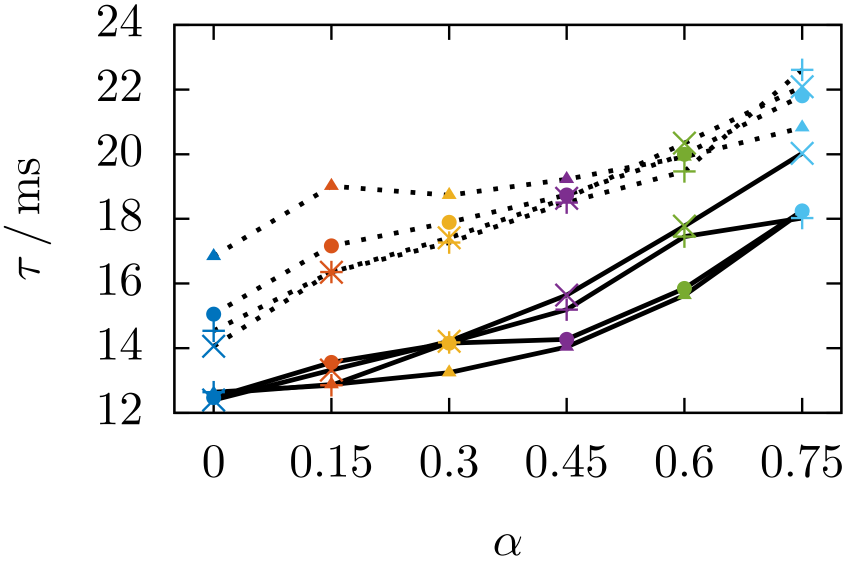

### S-9. CELL STIFFNESS VARIATION

All simulations in the manuscript were performed using a cell with a fixed shear modulus. We plot in figure S-8 the results of figure 9 and figure 10 (gray lines) together with the same data of a softer cell with a shear modulus of $\mu = 500\,\mathrm{Pa}$.

The maximum cell stress in figure S-8(a) to (e), i. e., the peak right after the exit as well as the magnitude inside the nozzle channel before the exit, are approximately half of the value obtained for the stiffer cell. This is due to the stress calculation in (9), where the shear modulus scales the influence of the deformation. Additionally, the cell strain peaks in figure S-8(f) and (g) are of similar order. An inverse scaling with the stiffness is observed for the stress relaxation time $\tau$ in figure S-8(e), showing values about twice as large for the soft cell compared to the stiff one.

FIG. S-8. Influence of the cell stiffness: (a) to (e) data from figure 9 and (f, g) data from figure 10 for a softer cell with $\mu = 500\,\mathrm{Pa}$.

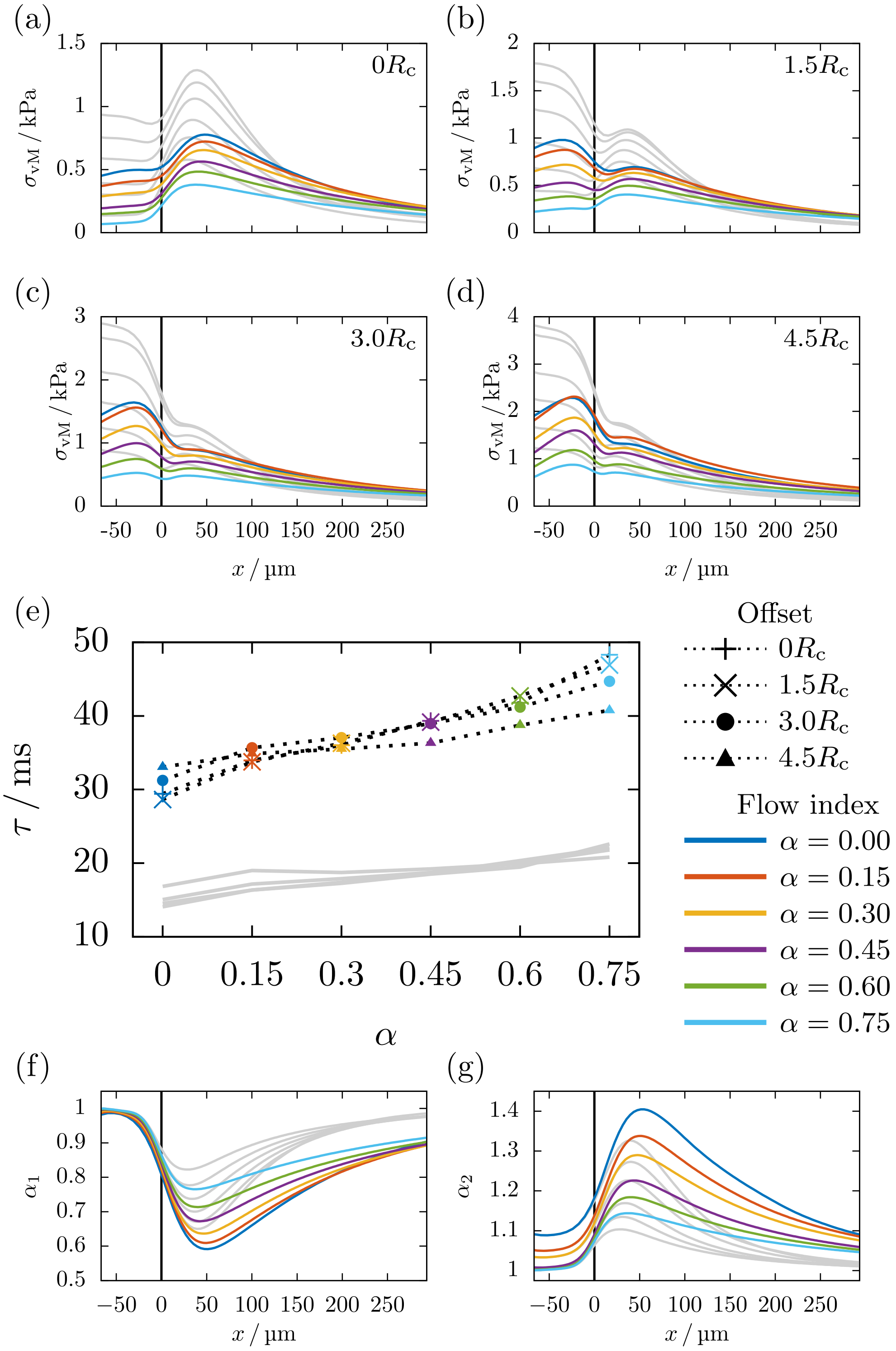

FIG. S-9. Estimated elongational stress at the nozzle exit for a bioink with shear thinning exponent $\alpha = 0.6$ in differently sized nozzles and with (a) variable extrusion speed $u_{\mathrm{avg}}$, or (b) different flow rate $\Omega$ or (c) different printing pressures.

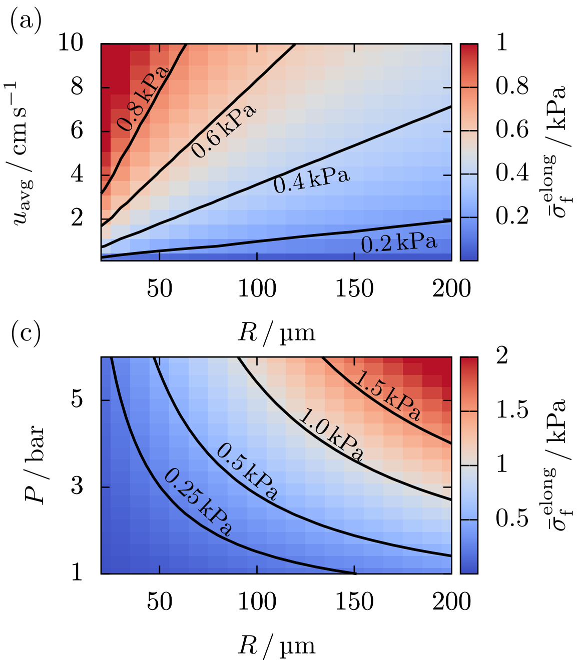


### S-10. ELONGATIONAL FLOW ESTIMATE

To compute the average elongational stress $\bar{\sigma}_{\mathrm{f}}^{\mathrm{elong}}$ acting on cells right at the nozzle exit, we start by calculating the maximum and average flow velocity of our bioink using our tool from [5], as in figure 1(b). From equations (15) and (16) we then obtain $\bar{\sigma}_{\mathrm{f}}^{\mathrm{elong}}$ as a function of the average extrusion velocity $u_{\mathrm{avg}}$ and the nozzle radius $R$. The result is shown in figure S-9, using a the same parameter space as in figure 13 of the manuscript.

[7] E. Guazzelli, J. F. Morris, and S. Pic, *A physical introduction to suspension dynamics*, Cambridge texts in applied mathematics (Cambridge University Press, Cambridge ; New York, 2012).
